# Artificial seawater based long-term culture of colonial ascidians

**DOI:** 10.1101/2021.01.31.429038

**Authors:** M. Wawrzyniak, L.A. Matas Serrato, S. Blanchoud

## Abstract

Tunicates are highly diverse marine invertebrate filter-feeders that are vertebrates’ closest relatives. These organisms, despite a drastically different body plan during their adulthood, have a tissue complexity related to that of vertebrates. Ascidians, which compose most of the Tunicata, are benthic sessile hermaphrodites that reproduce sexually through a motile tadpole larval stage. Over half of the known ascidians species are able to reproduce asexually by budding, typically leading to the formation of colonies where animals, called zooids, are interconnected through an external vascular system. In addition, colonial ascidians are established models for important biological processes including allorecognition, immunobiology, aging, angiogenesis and whole-body regeneration. However, the current paucity in breeding infrastructures limits the study of these animals to coastal regions.

To promote a wider scientific spreading and popularity of colonial ascidians, we have developed a flexible recirculating husbandry setup for their long-term in-lab culture. Our system is inspired both by the flow-through aquariums used by coastal ascidian labs, as well as by the recirculating in-lab systems used for zebrafish research. Our hybrid system thus combines colony breeding, water filtering and food culturing in a semi-automated system where specimens develop on hanging microscopy glass slides. Temperature, light/dark cycles, flow speed and feeding rates can be controlled independently in four different breeding environments to provide room for species-specific optimization as well as for running experiments. This setup is complemented with a quarantine for the acclimatization of wild isolates.

Herein we present our success in breeding *Botrylloides diegensis*, a species of colonial ascidians, for more than 3 years in recirculating artificial seawater over 600 km away from their natural habitat. We show that colonies adapt well to in-lab culturing provided that a suitable marine microbiome is present, and that a specific strain can be isolated, propagated and efficiently used for research over prolonged periods of time. The flexible and modular structure of our system can be scaled and adapted to the needs of specific species, such as *Botryllus schlosseri*, as well as of particular laboratory spaces. Overall, we show that *Botrylloides diegensis* can be proficiently bred in-land and suggest that our results can be extended to other species of colonial ascidians to promote research on these fascinating animals.

**Highlights:**

- First artificial seawater based recirculating aquaculture for colonial ascidians
- Over 3 years of continuous breeding
- Semi-automated setup with minimized maintenance
- Good biomass production for strain propagation
- 4 different culture conditions for optimized breeding for species of interest

## 1. Introduction

Tunicates are a group of over 3000 known extent highly diverse worldwide marine invertebrates, the majority of which are sub-tidal suspension-feeding hermaphrodites (Holland, 2016). Tunicata is separated into three classes: Appendicularia, Thaliacea and the paraphyletic Ascidiacea itself composed of the three orders Phlebobranchia, Aplousobranchia and Stolidobranchia (Delsuc et al., 2018; Kocot et al., 2018). Appendicularia has 68 identified species of planktonic free-swimming organisms that possess chordate traits common to most tunicate larvae including a notochord, neural tube and pharyngeal gill slits (Holland, 2016). These animals form communities where each individual is enclosed inside a special external mucous structure, termed house, which concentrates and funnels their food (Lambert, 2005). Thaliacea is composed of 77 species of planktonic pelagic animals forming free-floating compound colonies (Lambert, 2005). These organisms alternate between sexual reproduction to initiate new colonies, and asexual reproduction by budding to increase the size of the colony. Ascidians are defined morphologically as sessile benthic organisms, while phylogenomic analyses have shown that the pelagic Thaliaceans are nested within this paraphyletic class (Delsuc et al., 2018; Kocot et al., 2018). With 2935 currently identified species, of which 1730 are colonial species (Shenkar and Swalla, 2011), the paraphyletic taxon Ascidiacea is by far the most diverse class of tunicates. Ascidians are present in various marine habitats, from sandy beds to rocky shores and from coral reefs to kelp forests (Shenkar and Swalla, 2011). In addition, ascidians very efficiently colonize artificial substrata and have thus become a major source of biofouling, with a negative impact on economical sectors such as shellfish farming and marine transport (Aldred and Clare, 2014; Comeau et al., 2015).

However, ascidians also represent great opportunities for research as well as for the pharmaceutical and biomedical industry. First, Tunicata’s phylogenetic position as vertebrates’ closest relatives (Delsuc et al., 2006) renders them key models for evolutionary research, in particular to study the emergence of chordate-specific processes. Second, colonial ascidians are the only known chordates capable of undergoing whole-body regeneration (WBR) whereby a fully functional adult is restored from a minute fragment of vasculature in as little as 10 days (Brown et al., 2009; Rinkevich et al., 1995; Voskoboynik et al., 2007). Third, their recently established application as bioindicators of micro-pollution (Navon et al., 2020; Tzafriri-Milo et al., 2019; Vered et al., 2019) combined with their ubiquitous localization makes these animals a perfect tool for environmental bio-surveillance. Fourth, some colonial ascidians have emerged as model organisms for an increasing number of research fields, including immunobiology, allorecognition, angiogenesis and aging (Ballarin, 2008; Gasparini et al., 2015a; Kassmer et al., 2016; Munday et al., 2015; Paz and Rinkevich, 2002). In addition, ascidians are a particularly promising source of marine natural products (MNP) with already over 1000 isolated molecules (Chen et al., 2019) and numerous reported applications including antibacterial, anti-fungal, anti-viral, anti-inflammatory and antitumoral (Palanisamy et al., 2017). There is thus a tremendous scientific interest in the study of ascidians.

In our lab, we are particularly interested in *Botrylloides diegensis* (**Fig. 1A**), an invasive species widely present worldwide but commonly misidentified as *Botrylloides leachii* (Bay-Nouailhat and Bay-Nouailhat, 2020; Viard et al., 2019), that we collect in the Northern Mediterranean sea (**Fig. 1B**). *B. diegensis* is the first and currently only tunicate species in which a population of stem-cells has been shown to be responsible for its capacity to undergo WBR (Kassmer et al., 2020). Moreover, the recently identified erroneous assignation of *B. diegensis* genotyping reference sequences (Viard et al., 2019) to the well studied species *B. leachii* (Blanchoud et al., 2017, 2018b; Brunetti, 1976; Cima et al., 2001; Paz and Rinkevich, 2002; Rinkevich et al., 1995, 2007, 2010; Zondag et al., 2019, 2016) suggests that *B. diegensis* might be more widely studied than reported.

**Figure 1.**
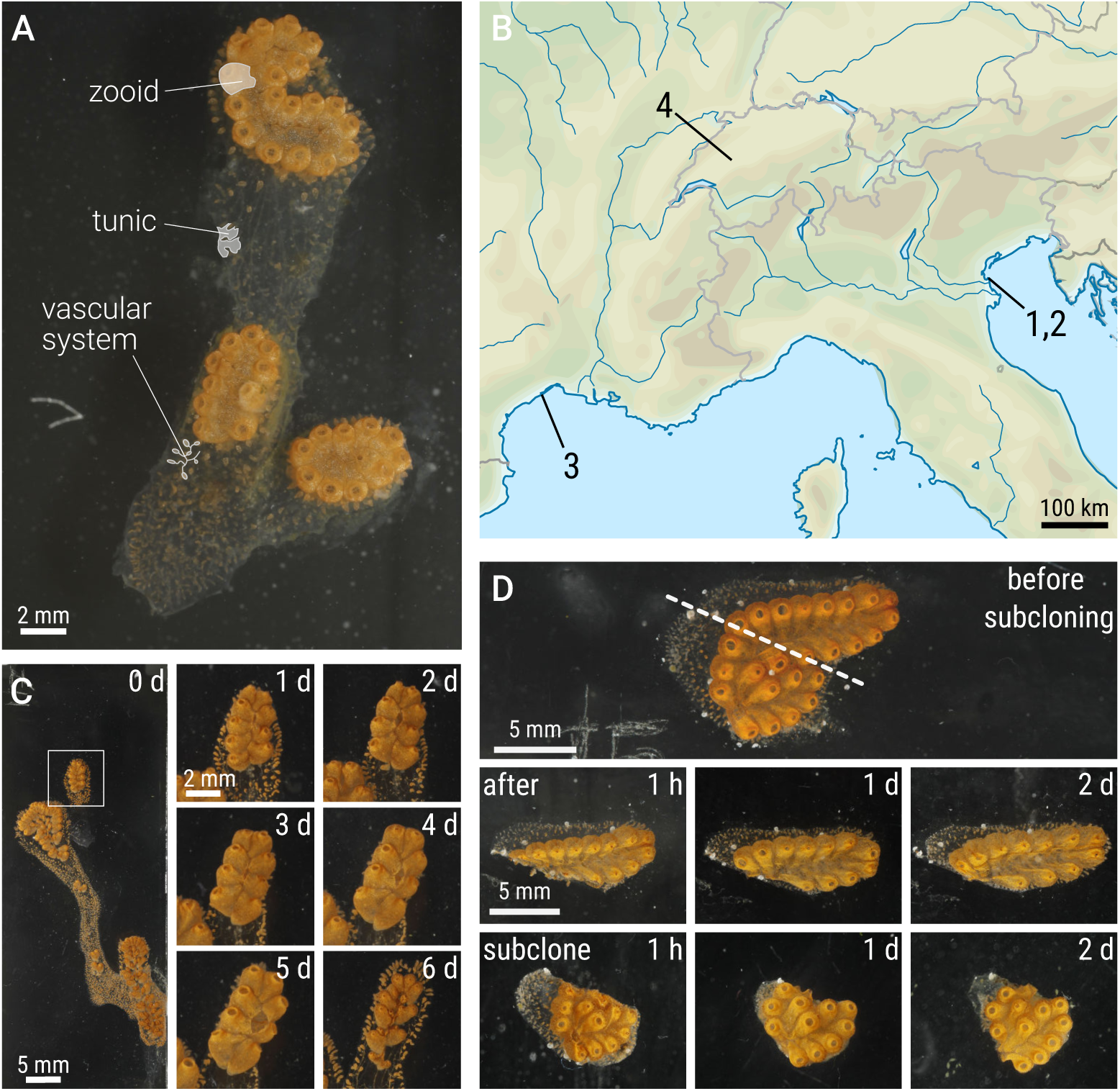
Breeding *Botrylloides diegensis* in artificial seawater. (**A**) A typical colony of *Botrylloides diegensis* settled on a glass slide. Representative portions of the indicated parts of the colony are highlighted by a light gray shading. (**B**) The sampling sites where colonies were collected (Chioggia, IT: 1, 2; Sète, FR: 3) and location of the husbandry system (Fribourg, CH: 4). (**C**) Asexual reproduction cycle in our breeding system. The colony growing on slide 117 is shown as an example of the observed weekly blastogenic cycle (days 0 to 6). Blastogenesis starts and ends with the takeover stage, where the adult filter-feeding zooids are being resorbed by the colony. White square highlights the system that is magnified and followed every day throughout the week, overlaid with the corresponding number of days after takeover. (**D**) Example of subcloning a colony. The colony growing on slide 91 is shown as an example of subcloning with its lower half being moved to a second slide. Shown are the status of both fragments at the indicated time post surgery, which is depicted as a dashed line overlaying the intact colony.

*Botrylloides diegensis* colonies are composed of numerous genetically identical adults, known as zooids, embedded in a gelatinous matrix, called tunic, that contains a shared external vascular system (**Fig. 1A**). Individual zooids are approximately 2–3 mm in length and have a complex morphology with organs homologous to that of vertebrates, including gill slits, a gastrointestinal tract, a nervous system, endocrine glands and a heart (Brunetti, 2009). *Botrylloides diegensis* life cycle begins seasonally (Rinkevich et al., 1993) from a tadpole larval stage with a typical chordate body plan (Berrill, 1947). Following a free-swimming phase, the larva settles and metamorphoses into a sessile adult known as oozooid. This founding animal will then expand into a colony through cycles of asexual reproduction termed blastogenesis whereby up to three new animals, known as buds, are produced by each zooid (**Fig. 1C**; (Berrill, 1947). Under optimal environmental conditions, the resulting clonal colony is thus an exponentially growing and highly modular structure (Hiebert et al., 2020; Marfenin, 1997). These features, combined with a great healing capacity, allows researchers to separate large colonies into smaller clonal replicates, called subclones, that will reproduce and propagate independently (**Fig. 1D**; (Manni et al., 2007). These advantages make colonial tunicates, and *Botrylloides diegensis* in particular, highly attractive and pertinent marine model organisms.

Ascidian research is an active community with numerous groups working worldwide using three main culturing methods. One most straightforward technique is onsite processing (Roberts et al., 2007; Sheets et al., 2016; Zondag et al., 2016; Ueki et al., 2018; Hiebert et al., 2019). No long-term culture is involved, animals are sampled in their natural environment and directly processed for downstream analysis. This approach is thus well suited for studying *in situ* physiological aspects of Tunicates but it requires a reasonably close-by sampling site and can be influenced by the potential seasonal availability of given species in some regions of the globe (Coma et al., 2000). A second prominent approach is the culture in a flow-through system (Hirose et al., 1995; Taketa et al., 2015; Blanchoud et al., 2017; Scelzo et al., 2019), typically with filtered natural seawater from the site of sampling, which provides close to natural environmental conditions including in terms of feeding diet. This method allows to isolate and maintain strains on the long-term in confined and hazard-free conditions. It does require either a close proximity to a source of seawater or a consequent organization to provide for the necessary amounts of seawater. A third common approach is the culture in a closed system with periodic water changes (Brown et al., 2009; Epelbaum, 2009; Rinkevich et al., 2010). This setup is most typically used with water from the sampling site for short-term culture of animals, although it has been used for extensively long studies too (Rinkevich and Shapira, 1998), as well as with artificial seawater (Nagar and Shenkar, 2016). The main limitations of this approach is the maintenance it requires on the long term, and the amount of water it depends upon when culturing a large number of colonies simultaneously. Consequently, all the labs that have the opportunity to do so combine these three approaches to benefit from their specific advantages. Yet, it consistently requires a routine access to either wild specimens or natural seawater and thus restricts their application to coastal locations. The only exception to this spatial limitation that we are aware of is that of Joly et al. who developed a recirculating system for breeding transgenic lines of the solitary ascidian *Ciona intestinalis* (Joly et al., 2007). Consequently, there is currently, to our knowledge, no recirculating husbandry system for the culture of colonial ascidians, which spatially limits their study and thus hinders their popularity as model organisms.

Here, we present for the first time a unique husbandry system designed for the in-land breeding of sessile colonial tunicates based on recirculating artificial seawater (**Fig. 2-3**). We have been successfully using this setup for over 3 years with the continuous presence of some of our strains, combined with a periodic addition of new wild isolates. Our recirculating system runs on artificial seawater and uses commercial algae pastes as feeding diets. Our aquarium is designed to permit up to 4 different culture conditions in parallel to optimize the breeding of the species of interest. In this study, we have been growing both *Botrylloides diegensis* as well as *Botryllus schlosseri* on microscopy glass slides. We present our developed aquaculture system, the maintenance setup used for the colonies, our long-term results in the clonal expansion of colonial ascidians, an example of a comparative breeding experiment and the importance of establishing a suitable marine bacterial community for the animals’ survival. Our data shows that we are able on the long term to abolish any spatial constrains with respect to the natural environment of colonial ascidians. This successful setup will enable research on colonial tunicates to be undertaken in novel places and will thus promote as well as popularize these fascinating animals as model organisms.

**Figure 2.**
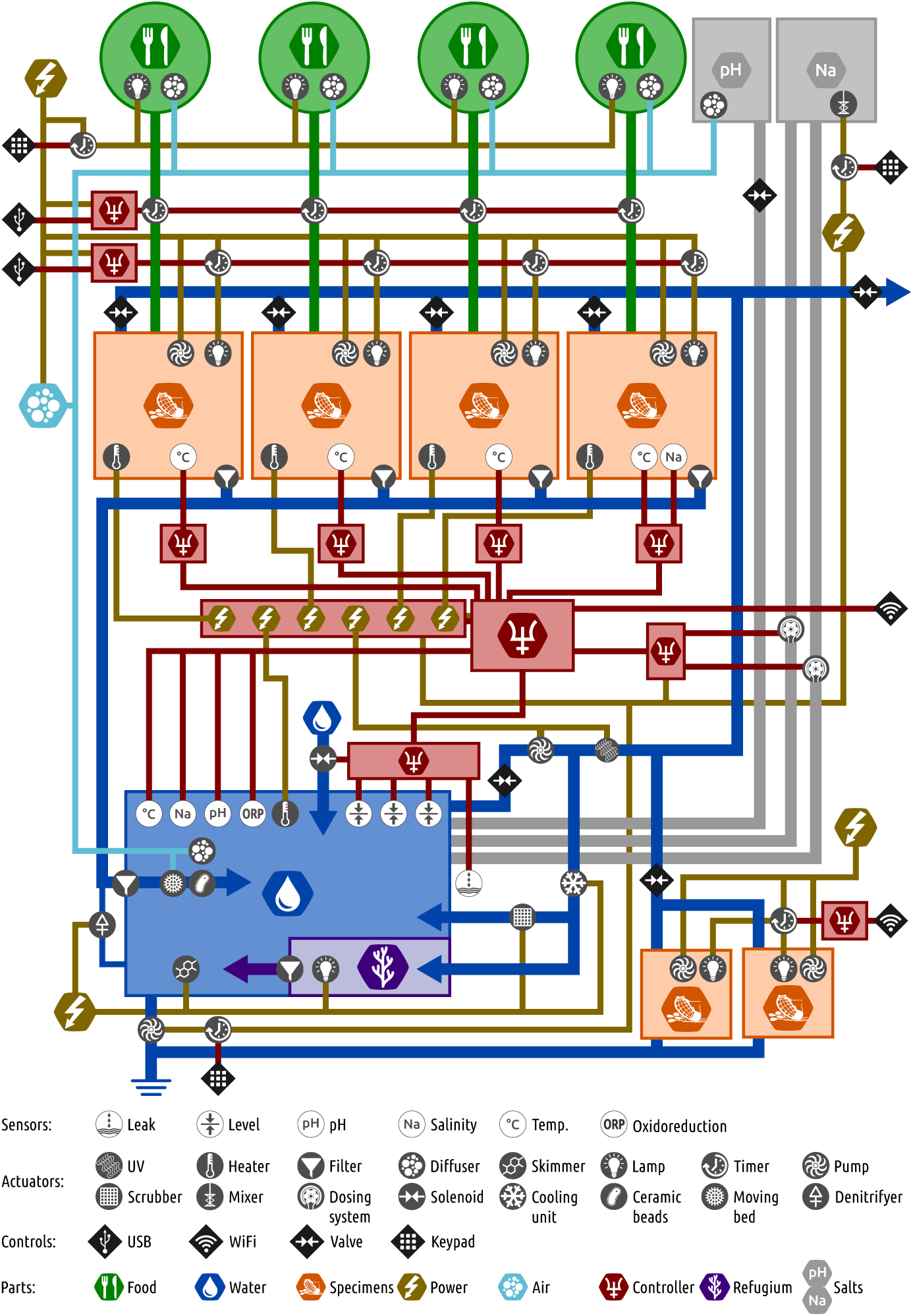
Schematic blueprint of our recirculating artificial seawater system. The components, their connections and control, as well as the conceptual structure of our husbandry system are depicted using the specified icons and color-codes. The two smaller specimens tanks visible in the lower right corner of the schema represent the two quarantine tanks.

**Figure 3.**
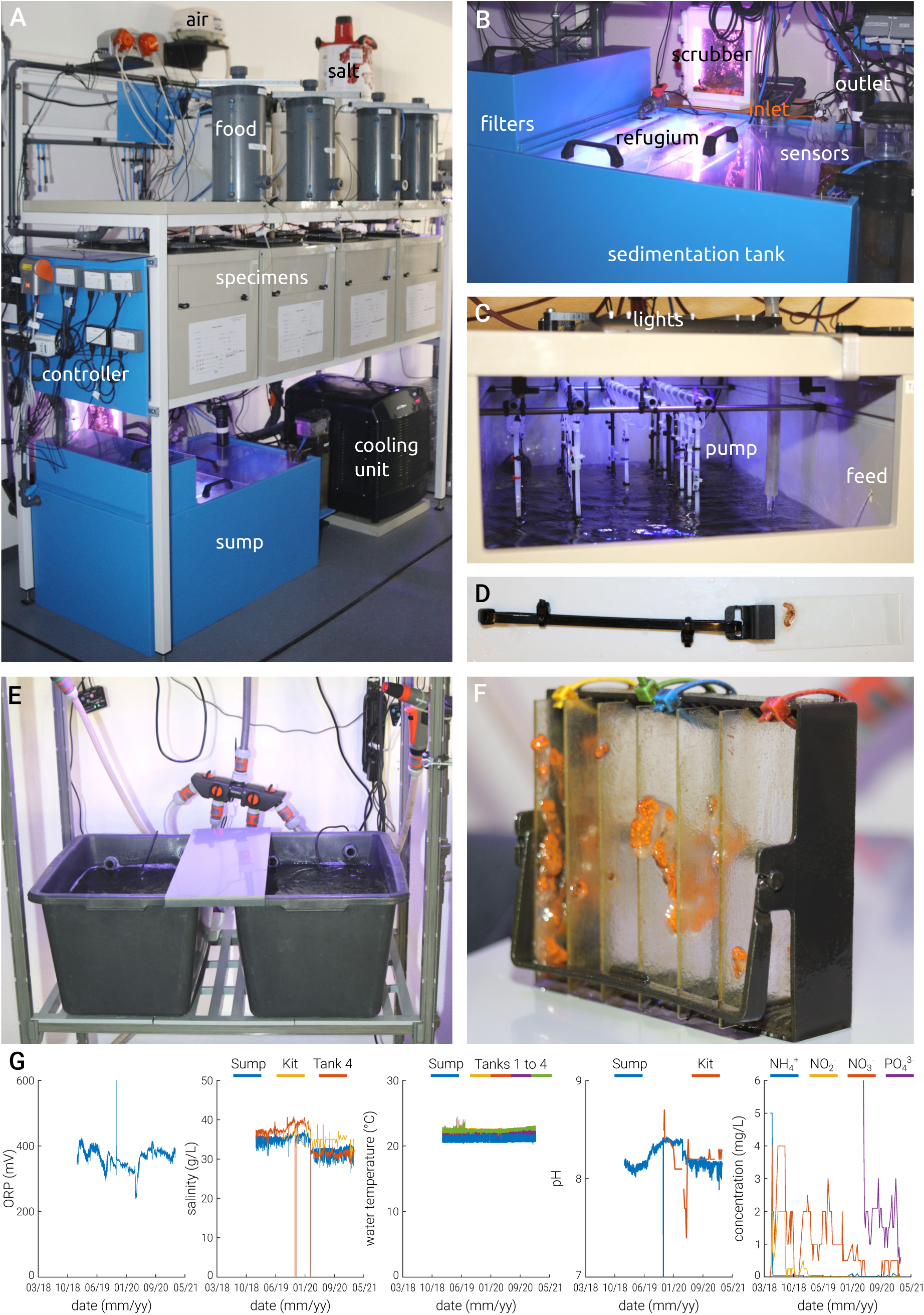
The colonial ascidian husbandry setup. (**A**) Picture of the developed prototype. The main system is composed of a shelf of tanks for food flowing into the specimens’ tanks recycling through the sump (**B**). View inside the sump where water is filtered and rectified automatically. (**C**) Vertical culture of colonial ascidians – each specimens’ tank is equipped with racks of hooks. (**D**) Colony hanger used to hold the glass slides vertically. (**E**) Two quarantine containers. (**F**) Staining racks are used during quarantine. (**G**) Parameters of the water quality measured in our system between March 2018 and March 2021 (**Table 2**). From left to right: ORP – oxidation reduction potential (mV); salinity (g/l); water temperature in the sump and tanks 1 – 4 (°C); pH in the sump and measured with a kit; levels of noxious substances: NH_4_, NO_2_, NO_3_ and PO_4_ (mg/l).

## 2. Materials and Methods

### 2.1 Aquarium structure

The complete list of material and equipment used to assemble our system is provided in **Table 1** (**Supplementary Figures S1-S4**). Briefly, the structure of the system is a SS 316 40×40 and 30×30 mm powder coated stainless steel stand (LxWxH 2.0 x 0.9 x 2.2 m), custom manufactured PE/PP 40 L specimen tanks, a custom manufactured PE/PP 200 L filtration sump as well as four custom manufactured PE/PP 10 L food culturing vessels. The system is equipped with the following materials: fountain pump (universal 300, Eheim, DE), air blower (Medo LA-60B, Nitto Kohki, DE), cooling unit (Titan 4000, Aqua Medic, DE), heaters (titanium D-D 600W, Schego, DE), motor driven protein skimmer (DC runner 1.1, AquaMedic Gmbh, DE), circulation pump (Titan-Tec 10000, Messner Gmbh, DE), UV steriliser (Vecton2, TMC Aquarium, US), and lights (AquaBeam 1500 Ocean blues, TMC Aquarium, US). The construction of our prototype was outsourced to a company specialized in aquatic research equipment (Zebcare, Fleuren & Nooijen BV, NL).

**Table 1.**
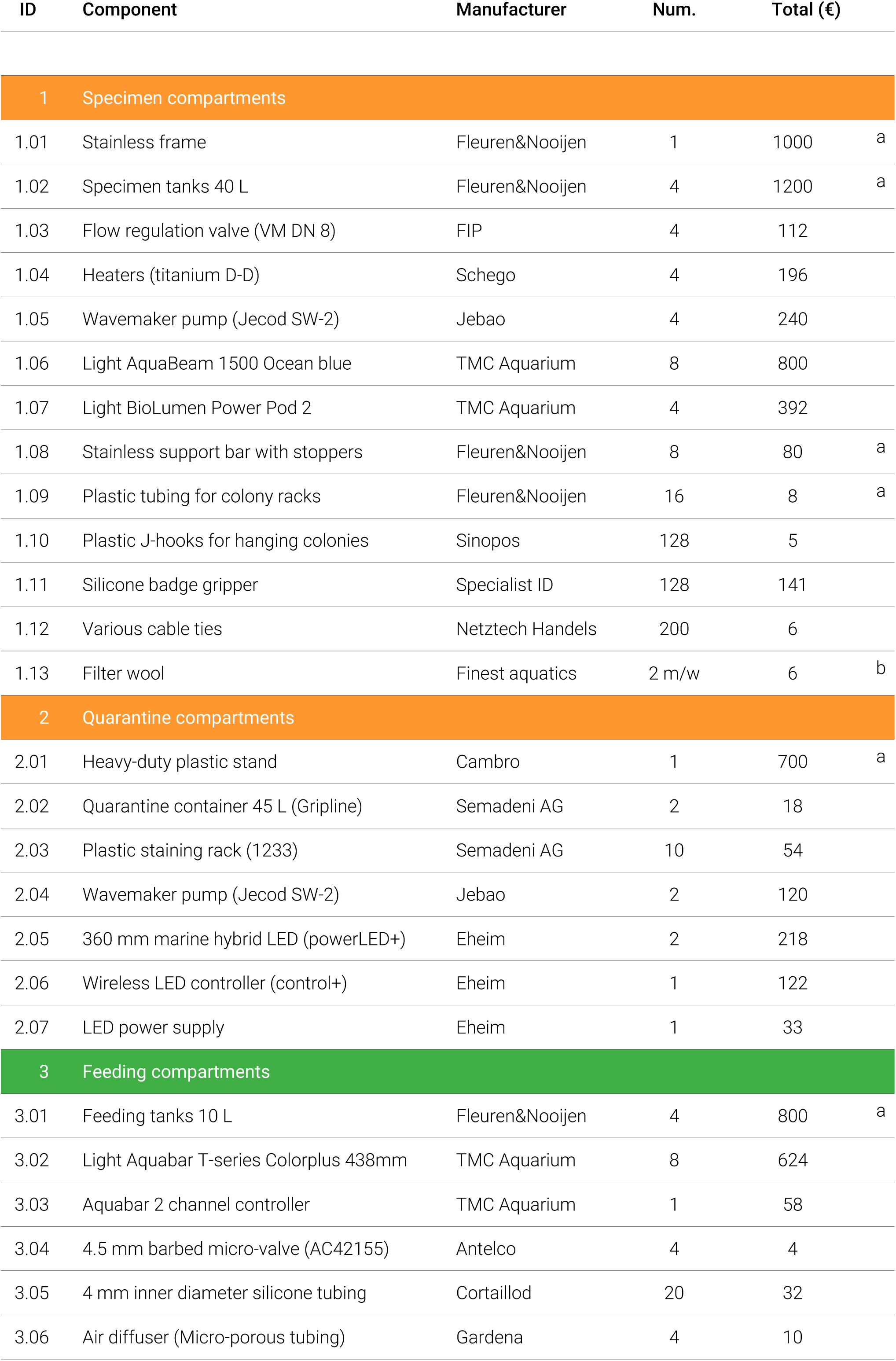
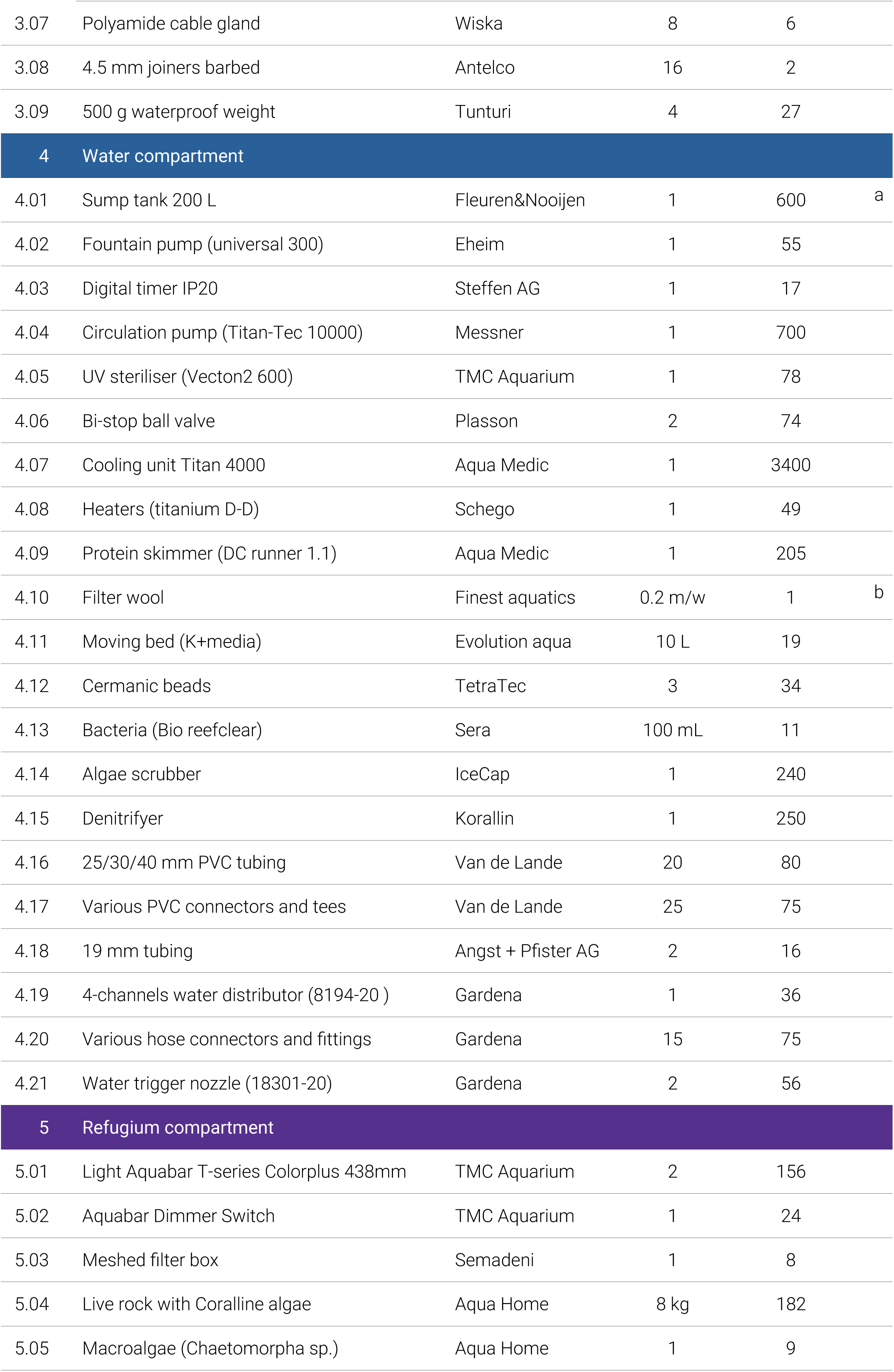
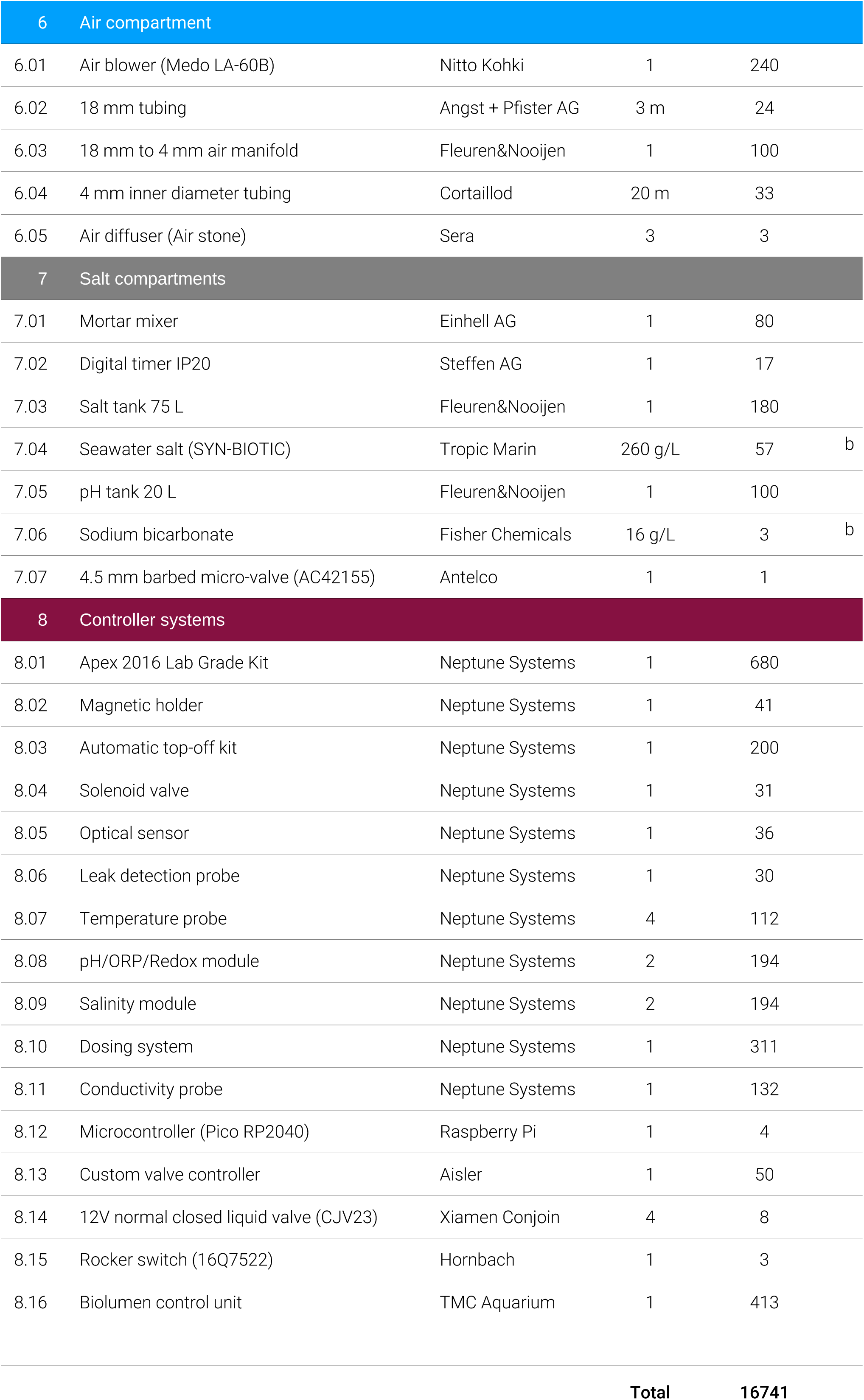
Components use in our husbandry system for colonial ascidians. Most components are visible in **Supplementary Figures S1-S4**.Note that the given prices have been gathered from a variety of different providers and currencies and will likely vary. Transportation, taxes and assembly costs were not taken into account. **a)** these components were custom-build and their costs are thus estimates only. **b)** these components are consumables and the given price corresponds to what is required for one complete setup.

A quarantine area composed of two flow-through 45 L tanks (Gripline, Semadeni AG, CH), aerated by one pump each (Wavemaker Jecod SW-2, Jebao, CN) and illuminated by a LED rack each (Marine hybrid powerLED+, Eheim, DE) was added to the main system by our departmental workshop. Our quarantine tanks are placed on a heavy-duty plastic stand (Camshelving Premium, Cambro, US). Staining plastic racks (1233, Semadei AG, CH) were used for hosting the animals on their glass slides.

The entire setup was equipped with a range of sensors to obtain a fully automatized control on the environment of the aquarium using the Apex (Apex System, Neptune Systems, US) controller, a custom microcontroller (Pico, Raspberry Pi, UK) as well as light controllers (BioLumen, TMC Aquarium, US). In particular, salinity, pH, day/night cycles, temperature and changes of water are performed automatically and reproducibly.

For the microbiome studies, two 10 L glass aquaria (NanoCube, Dennerle, DE) were used, filled with 7 L of artificial seawater (ASW), equipped with small pumps for water aeration (FP100, Sera, DE). Illumination using LED rack and animal hosting using staining plastic racks were identical to those used in our quarantine area. Three colonies were cultured together per glass aquaria for each microbiota condition, repeated with another set of three colonies for all conditions except inoculation with B001.

### 2.2 Artificial seawater production and filtration

All kits for water quality as well as Syn-biotic sea salt were purchased from Tropic Marin (DE). Seawater refractometer was purchased from Red Sea (US), Optic USB base station and Hobo Pendant temperature/light data logger were purchased from Onset Computer Corporation (US). Filtration is performed by a denitrifyer filter (Sulphur BioDenitrator, Korallin, DE), ceramic beads (Tetra, DE), nitrifying bacteria (Bio reefclear, Sera, DE), stirred plastic moving bed (K+Media, Evolution Aqua, UK), an algae scrubber (Medium Turf Scrubber, IceCap, US) and phosphate absorption resin (Brightwell Aquatics, US). Live stones cured with Coralline algae as well as macroalgae were bought from Aqua-Home (CH). Concentrated marine biota was made by sterile filtering 10 L of gravity filtered natural seawater using 0.22 µm Nalgene filters and resuspending the content of the filters in 50 mL of gravity filtered natural seawater. Other chemicals and small laboratory equipment were purchased from Thermo Fisher Scientific (US).

### 2.3 Animal breeding and handling

Numerous colonial ascidians (*Botrylloides diegensis* in particular) were collected in two locations in the coastal town of Chioggia (IT, 45.21306, 12.27394 and 45.21787, 12.28966, October 2017, May 2018, October 2018, May 2019) as well as in Sète (FR, 43.39608, 3.69889, March 2018). Most colonies were brought back to the lab with their original substrate in closed boxes filled with seawater, some were transferred to microscopy glass slides first and left to settle for 48 h before being transferred to the lab, but we observed a lower success rate with this second approach. Half a dozen mussels were also collected in Chiogga during our first visit to inoculate the husbandry system’s microbiota following a protocol developed for echinoderm breeding in artificial seawater (Oliveri, personal communication, unpublished). Colonial ascidians were then acclimatized from their *in situ* environmental conditions to our breeding conditions (**Table 2**) in our quarantines and transferred to 76×26×1 mm microscopic slides (Marienfeld, DE). No colony was observed to be gravid at the time of transfer. After the animals had properly settled on the slide, they were transferred to the recirculating system where they developed vertically inside the specimen tanks using a custom attachment setup using cable ties (Netztech Handels AG, CH) and a silicone badge gripper (Specialist ID, US). The genus of the colonies was determined visually based on the morphology of the systems, the species was determined for the 4 most proliferative species by DNA barcoding using the mitochondrial cytochrome oxidase I (mtCOI) gene, as described below.

**Table 2.**
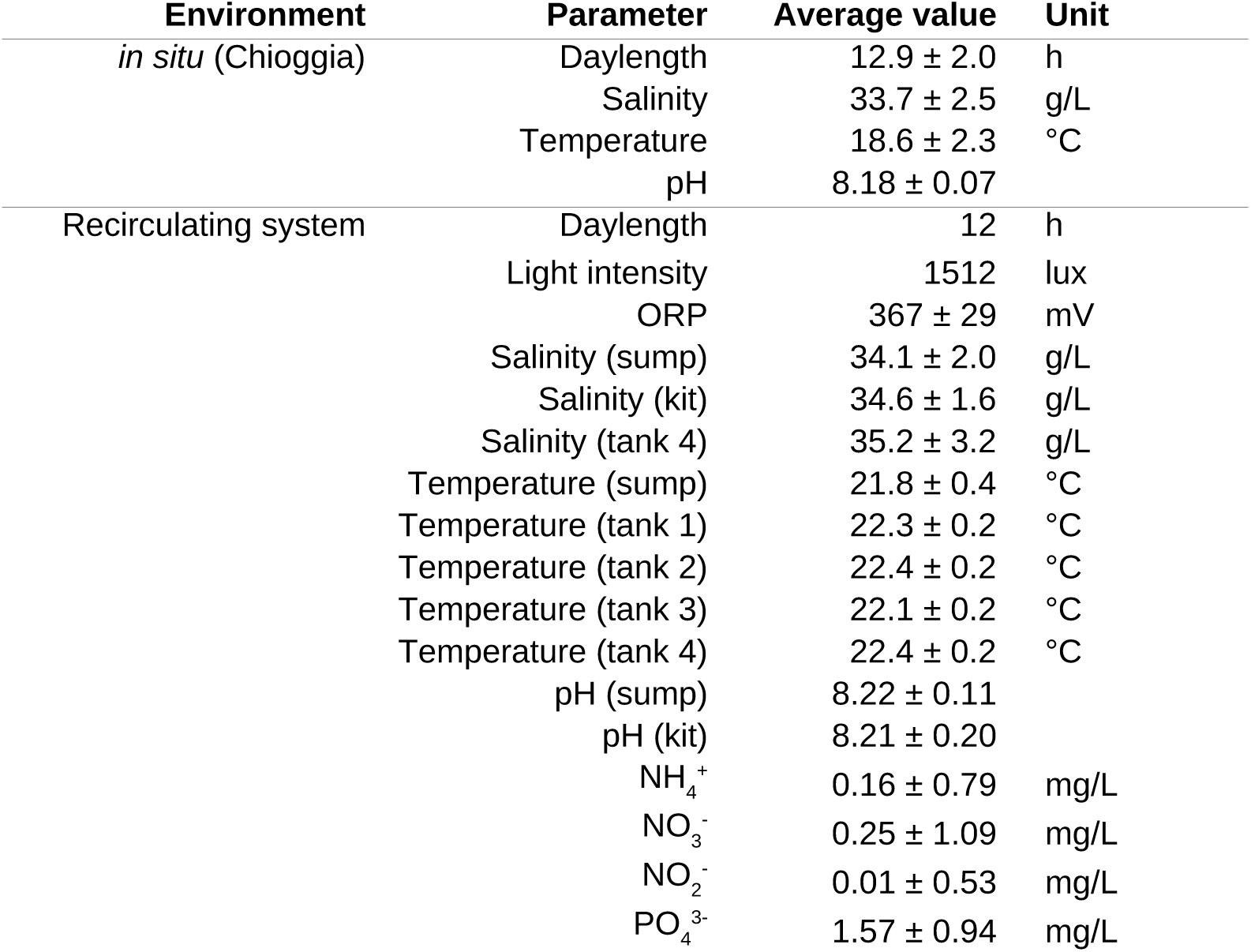
Comparison of in situ and breeding environmental conditions. Average and standard deviation of the various parameters measured in each environment.

| Environment | Parameter | Average value | Unit |
| --- | --- | --- | --- |
| *in situ* (Chioggia) | Daylength | 12.9 ± 2.0 | h |
|  | Salinity | 33.7 ± 2.5 | g/L |
|  | Temperature | 18.6 ± 2.3 | °C |
|  | pH | 8.18 ± 0.07 |  |
| Recirculating system | Daylength | 12 | h |
|  | Light intensity | 1512 | lux |
|  | ORP | 367 ± 29 | mV |
|  | Salinity (sump) | 34.1 ± 2.0 | g/L |
|  | Salinity (kit) | 34.6 ± 1.6 | g/L |
|  | Salinity (tank 4) | 35.2 ± 3.2 | g/L |
|  | Temperature (sump) | 21.8 ± 0.4 | °C |
|  | Temperature (tank 1) | 22.3 ± 0.2 | °C |
|  | Temperature (tank 2) | 22.4 ± 0.2 | °C |
|  | Temperature (tank 3) | 22.1 ± 0.2 | °C |
|  | Temperature (tank 4) | 22.4 ± 0.2 | °C |
|  | pH (sump) | 8.22 ± 0.11 |  |
|  | pH (kit) | 8.21 ± 0.20 |  |
|  | NH<sub>4</sub><sup>+</sup> | 0.16 ± 0.79 | mg/L |
|  | NO<sub>3</sub><sup>-</sup> | 0.25 ± 1.09 | mg/L |
|  | NO<sub>2</sub><sup>-</sup> | 0.01 ± 0.53 | mg/L |
|  | PO<sub>4</sub><sup>3-</sup> | 1.57 ± 0.94 | mg/L |

Taxonomically identified specimens of *Botryllus schlosseri* and *Ciona robusta* were obtained through the European Marine Biological Resource Center (EMBRC) network. *B. schlosseri* colonies were cultured together with the *B. diegensis* colonies using the same technique and environmental conditions, both on the larger 76×52×1 mm slides they were shipped onto, as well as on standard microscopic slides after they were subcloned. *C. robusta* animals failed to reattach to any new substrate once they were detached from the plastic Petri dish they were shipped onto. The *C. robusta* specimen was attached to a closed cable tie using a 4-0 prolene suture monofilament (Ethicon, Johnson & Johnson, US) threaded through its posterior tunic and hung with its oral siphon facing downwards on the hanger system.

Animals were fed using a blend of commercial algae concentrates from Instant algae (LPB Shellfish Diet, Shellfish Diet 1800, RG complete, RotiGrow Plus, US), complemented with a planktonic substitute based on crustaceans’ eggs extract (Zootonic, Tropic Marin, DE) and commercial baker’s yeast (Rinkevich and Shapira, 1998). Feeding in the recirculating system was controlled by 12 V normal closed solenoid valves (CJV23, Conjoin, CN) controlled by a customized microcontroller (Raspberry Pi Pico, UK, Supplementary File 1). Feeding of the quarantines and of the glass aquaria was done manually daily with 1 mL and 20 µL of the feeding mix, respectively. Colonies were monitored by macrophotography using a Canon 5D Mark II with a Sigma 50 mm f/2.8 EX DG macro objective and a Canon Speedlite 430EX II flash. Time-lapse was performed using a custom variant of FlyPi (Chagas et al., 2017) running on a Raspberry Pi 4B equipped with a Pi HQ Camera and a Pi 16 mm telephoto lens, imaging colonies placed in a Petri dish with flow-through water at a frame rate of one picture every 15 minutes.

Live L-type rotifers (Planktovie, FR) were purchased in brackish 15 g/L seawater. We thus acclimatized them to our system’s 35 g/L by slowly increasing the salinity of their medium by 5 g/L every week. Rotifers were cultured in 1 L Erlenmeyer flasks without agitation, fed with RG Complete every day and their water was changed every 2 days. Once the population became sufficiently dense, typically every other day, they were split in half. One half serving for continuing their expansion, the other half being fed to the specimen tanks.

### 2.4 Data collection and image analysis

Environmental parameters were collected from the daily measurements performed by the Hydrobiological Station of Chioggia (University of Padova, IT) available at https://chioggia.biologia.unipd.it/en/the-database/parameters-of-lagoon/. Daylength durations were collected from the reference website Time and Date AS (Stavanger, NO), for the location of Chioggia, available at https://www.timeanddate.com/sun/@12172566. Environmental parameters were gathered for the entire months during which collections took place in Chioggia (October 2017, May 2018, October 2018, May 2019), and their average and standard deviation calculated (**Table 2**).

Image analysis was performed manually on the weekly top-view pictures taken using our macrophotography setup. Total number of zooids present on both sides of each microscopy glass slide participating in the experiment was counted using Fiji (Schindelin et al., 2012) and the GNU Octave programming language (Eaton, 2012) with custom scripts was used to process the data. Zooids were identified by their oral siphon, marked using the multi-point region of interest (ROI) tool and organized using the ROI manager functionality of Fiji as compressed “.roi” files. For colonies undergoing takeover, resorbing zooids were counted until they became smaller than their buds.

### 2.5 Genotyping and bacterial culture

Colonies used for genotyping were cleaned using a small paintbrush and left 30 min to wash in ASW. Colonies were then briefly washed in clean ASW and 3-4 zooids were isolated using a single-edge razor blade. Genomic DNA was extracted using the HotSHOT method (Truett et al., 2000), its concentration measured using a NanoDrop 2000 (Thermo Fisher Scientific, US) spectrophotometer and the sample was diluted to reach 100 ng/µL using ultrapure water. PCR amplification was performed using 12.5 µL of MyTaq HS Mix (Bioline, US) hot-start polymerase, 2 µL of 10 mM forward primer, 2 µL of 10 mM reverse primer, 1 µL of 100 ng/µL genomic DNA and 7.5 µL of ultrapure water. The sequence of the published universal LCO1490/HC0219859 mtCOI primer pair was used (Folmer et al., 1994). PCR followed this program: 95 °C for 2 min, 35× (95 °C for 30 s, 50.5 °C for 30 s, 72 °C for 45 s), 72 °C for 10 min. PCR products were sent without purification for sequencing to Microsynth AG (CH). Species were assigned to each sequence by comparison with the NCBI NR database, considering the samples submitted in the context of (Viard et al., 2019).

Water samples were collected in sterile tubes, 100 µL were plated on 5 % agar lysogeny broth (LB) plates, spread using sterile glass plating beads and left to incubate at 37 °C for one week. Plates were then imaged using a Panasonic Lumix DMC-FZ150 camera, different bacterial types were identified visually and their respective concentration estimated by manual counting. Representative colonies were picked and amplified in 2 mL liquid LB culture at 37 °C overnight, from which 100 µL were plated again on LB plates. Single colonies were picked from the LB plates and further amplified in 2 mL LB for one week. 50 µL of the amplified bacteria (corresponding final concentrations of B001 2*10^-3^ bacteria per mL, B004 530 bacteria per mL and B005 5200 bacteria per mL, respectively) were used to inoculate freshly prepared ASW for the microbiome experiments, while another 500 µL were diluted and 100 µL of each dilution were plated on LB agar plates to estimate the corresponding bacterial concentration by manual counting. Genotyping was performed on isolated bacteria colonies picked on the LB agar plate or on the original agar plate for bacteria that did not propagate. Genomic DNA was extracted by lysing bacteria in 100 °C distilled water for 10 min. PCR amplification was performed using 12.5 µL of Accustart II PCR toughmix (Quantabio, US) polymerase, 0.5 µL of 25 mM forward primer, 0.5 µL of 25 mM reverse primer, 0.5 µL of loading buffer, 5 µL of genomic DNA and 6 µL of distilled water. The sequence of the published universal 27F/1492R 16S primer pair was used (Lane, 1991). PCR followed this program: 94 °C for 3 min, 35× (94 °C for 30 s, 56 °C for 30 s, 72 °C for 100 s), 72 °C for 10 min. PCR products were purified using the E.Z.N.A. Cycle Pure Kit (Omega Bio-tek, UK) and sent for sequencing to Microsynth AG (CH). Species assignation was done by performing a Megablast search of the sequencing results onto the NR database.

All sequences are available in **Supplementary File 2**.

## 3. Results

### 3.1. Description of the system

We designed a custom recirculating system for the in-land culture of colonial tunicates (**Fig. 2**), which allows to reduce periodic water changes to minimal amounts. In such closed systems, the wastes produced by the animals (e.g. NH_4_, NO_2_, NO_3_) accumulate over time and need to be continuously purified. Our modular aquaculture system is structured in three levels around a stainless stand to maximize the use of gravity for the circulation of the artificial sea water (ASW) in the aquarium. On the top, food culturing tanks distribute their content into the specimen tanks located in the middle, which overflow into the water filtration units placed at the bottom of the apparatus (**Fig. 3A**). The filtration unit combines mechanical and biological filters to reach a minimal weekly water change of 5 % of the total 450 L of ASW present in our system. Filtration includes a particle filter, a so-called moving bed filter (or upflow filter), a denitrifyer filter, an algae scrubber, a protein skimmer, live cured stones and a UV-C unit (**Fig. 3B**).

To breed our animals in optimal conditions, we specifically aimed at using a maximum of food-grade plastic material to avoid rust, wear and chemical contamination as much as possible. Four 40 L specimen tanks were custom made from 15/20mm PE/PP material. Tanks have one compartment and each has a heater which is placed at the bottom of the tank and is controlled by a temperature sensor placed on the side of the tank. Aeration and mixing of the water are ensured by a circulation pump placed on the back side of each tank. Tanks are completely closed to make a light-isolated environment, with a convenient front opening (**Fig. 3C**) for easy access and cleaning. Water falls into the tank through a silicone tube, which flow rate can be regulated to modulate the amount of flow-through, and leaves the tank from an outlet located in the back of the tank. A fleece is placed at the bottom of the tanks to collect solid wastes. Each tank is equipped with two lights sunken in the lid of the tank in such a way that the heat sink at the back of the LED tile is outside the tank for optimal heat transfer. The intensity and photoperiod of each LED unit can be controlled independently through a central control unit. Animals are placed in the tanks on a custom system of hangers made using plastic tubes, plastic hooks and badge grippers (**Fig. 3D**).

The central filtration unit (sump) has been placed on the ground (**Supplementary Figure S1)**. Water from the tanks is drained through a fleece filter into an aerated moving-bed biofilter, through a ceramic-beads biofilter and into the main sedimentation tank. The sedimentation tank is connected to an external protein skimmer with its submerged pump, a nitrate filter, an algae scrubber and a refugium populated with live cured stones and macro-algae. The refugium and the algae scrubber are constantly illuminated by two LED units. The sedimentation tank is connected to the main circulation pump, which provides system water to the cooling unit, the algae scrubber and the refugium in a closed loop, as well as to the specimen tanks through a UV sterilizer. In addition, periodic water changes are controlled by a timer-activated submerged fountain pump. This pump discards system water into the sewage through the sump’s overflow drain to induce its replacement with fresh seawater. This pump can also be used for complete drainage of the system. In order to reduce evaporation, the whole filtration tank is fitted with a clear plastic lid.

Above the specimen tanks, we have installed four 10 L cylindrical feeding tanks (**Supplementary Figure S2)**. Each tank is made of non-transparent PVC pipes 200 mm in diameter and 300 mm long. An external water level made of clear PVC hose and a visit window allow to monitor the content of each tank. Content of the tanks is mixed by injecting air through a microporous tube placed at the bottom of the tank, and lighting is done by two LED rails placed inside a clear pipe through the center of the non-air tight lid of the tank. Food is drained outside of each tank by gravity through an electronically controlled micro-valve placed close to the bottom of the tank (**Supplementary Figure S3)**. Air is supplied via 4 mm silicone hoses directly from the main air pump through a custom-built manifold (**Supplementary Figure S4)**.

Besides the main system, we also maintain two quarantine containers (**Fig. 3E**) for new comers and new subclones (i.e. colonies mechanically separated from a bigger one and placed onto a new glass slide; **Fig. 1D**). This part has been designed and built up on site and is on purpose not recirculating into the main system. However, both tanks can be filled with system water, and system water can be collected there for experiments using handy garden hose nozzles. Each container can be filled with up to 30 L of saltwater, aerated and mixed by a pump placed in the back of the tank. Each tank is illuminated with photoperiod-controlled lights. Animals are placed in standing plastic staining racks (**Fig. 3F**) inside the quarantine tanks.

The conditions in the system are monitored and logged automatically (**Fig. 3G, Table 2**) using an APEX system from Neptune Systems (**Supplementary Figure S3)**, connected to a number of sensors and actuators. Temperature, salinity, pH and ORP (oxidation reduction potential) are measured every 2 min in the main sedimentation tank. Temperature is controlled by the cooling unit and a heater controlled by Apex. Salinity can be increased by pumping 270 g/L brine water into the system from a 75 L tank placed behind the feeding tanks using two peristaltic pumps, also controlled by Apex. Brine tank is mixed four times per day using a heavy duty concrete mixer controlled by a timer (**Supplementary Figure S4)**. An alert email is sent if some values deviate from their defined ideal range (7.9-8.5 pH, 27-35 salt, 14-24 °C, 300-400 ORP). Water level is monitored using two optical sensors driving an automatic top-off kit that delivers deionized water directly in the main sedimentation tank by opening a solenoid valve connected to the University’s water system (**Supplementary Figure S1)**. Temperature is also measured individually in each specimen tank, and salinity is measured in one of the tanks for redundancy. Leaks and water loss are monitored using a floor sensor and an optical sensor placed at the bottom of the sedimentation tank, both leading to the shutdown of the whole system in case of emergency. Additional water quality parameters (pH, NH_4_, NO_2_, NO_3_, PO_4_) and the conditions in the quarantine are monitored manually.

### 3.2. Initialization and maintenance of the system

Before its first usage, the aquaculture system was cured over six weeks. First, the system was filled with tap water, run for 48 h and completely emptied. This wash was repeated two more times to remove all traces of chemicals from the system. During this time, the wild mussels were acclimatized and cleaned from parasites in our quarantine using ASW changed once per week and fed using 1 mL/day of Shellfish Diet 1800. The cleaned aquarium was then filled up with ASW and curing was started. Ammonium chloride was added (3 mg/L) together with the nitrifying bacteria, the concentrated marine biota, the mussels and the live rocks, and the system was left to run for 24 hours. 5 grams of commercial fish food mix was added in the sedimentation tank followed by 7 g every week. Water quality (pH, NH_4_, NO_2_, NO_3_) was measured every other day until ammonium concentration started to decrease (1.5 weeks post inoculation). 1.5 mg/L of ammonium was then added and the monitoring continued. When the nitrite concentration started to decrease (4 weeks post inoculation), the nitrate filter was started following the manufacturer’s instructions. 0.5 mg/L of ammonium was added and the system monitored until the water quality was within the range defined by commercial kits (6 weeks post inoculation in total). System was then used following our routine maintenance as described below.

Maintenance of our aquaculture system was reduced to a minimum by automatizing as many steps as possible, the bulk of the remaining manual labor being dedicated to tending to the various needs of the animals. Weekly duties include only the cleaning of the protein skimmer and every other week the replacement of the fleece placed at the bottom of the specimen tanks for trapping solid waste as well as the cleaning of the algae scrubber. Once a month, the excess of algae growing in the refugium is removed, the live stones scrubbed clean and the Apex system checked. Every 2-3 months depending on our use of system water, the brine tank needs refilling, and once a year a general maintenance is done where all the tubing pumping brine water are thoroughly cleaned.

In our hands, the most important factors for a reliable performance of our recirculation system is the quality of the water; not inasmuch as having perfectly low concentrations for all the noxious compounds in the system water, but in preventing these compounds from accumulating to dangerously high concentrations. Consequently, we manually measured on a weekly basis the four most important ions (NH_4_, NO_2_, NO_3_, PO_4_, **Fig. 3G**) as well as pH and salinity to confirm the readings from our controller. In addition, we tested monthly the following other parameters: KH, O_2_, I, Ca_2_, Mg_2_, Si and K^+^.

### 3.3. Breeding colonial tunicates in-land

Wild isolates were collected twice a year (Spring/Autumn) in the Venetian lagoon (**Fig. 1B**). Animals were transferred on their original substrate (most typically mussel shells, flat green algae, solitary ascidians or the dirt present on the side of the pontoon) into 1 L water-tight wide-mouth cylindrical containers. Animals were then transported to the laboratory, which took on average between 8 h and 10 h. Important parameters for successful transportation of the collected samples are first a large animal to volume ratio (> 20X), regular water changes during transportation (every 2-3 h), as little air in the container as possible and minimizing the turbulences during transport. However, the best predictor of a colony that will well tolerate being transferred was its size, namely medium-sized colonies composed of 50-100 zooids were typically fitter once in our laboratory. Another important aspect to consider when collecting is the type of substrate the animal is growing on, and its shape. Colonies growing flat on mussels were the most appropriate for later transfer to our glass slides, while those intermingled with brittle algae were very difficult to isolate back in the lab.

Upon arrival, wild specimens were transferred to 2 L tanks filled with ASW adapted to the salinity and pH of the natural sea water they were transported in (**Table 2**). Animals were cleaned, their substrate reduced to a minimum and the ones without any sign of filter-feeding activity discarded. Animals were placed in our quarantine tanks, fed twice daily with 1 mL of feeding mix and left 24 h to adapt. On the next day, colonies were subcloned to numbered microscopy glass slides using a single-edge razor blade following a customized protocol (Bugada and Blanchoud, 2021). Slides were placed in plastic staining racks at a density of 7 slides per rack standing at the bottom of the quarantine tank. Colonies were left for 48 h to attach to the slides, carefully cleaned (Blanchoud and Wawrzyniak, 2021), the decaying pieces of tissue discarded and the remaining attached colonies placed in the second quarantine tank. Water in the quarantines was changed when the NH_4_ levels rose above 1.5 mg/L, or at least once a week. We have observed that in our 30 L setup, with up to two dozen colonies, this will take around 7 days. The colonies that detached were subcloned again into the first quarantine and the process repeated until all colonies either attached or decayed. Colonies were then acclimatized to the environmental parameters of our recirculating system by gradually adapting the parameters of the quarantine over two weeks. The breeding environmental conditions were chosen to be as close as possible to the *in situ* environmental conditions of the colonies, although temperature was increased to reduce transitions with bench work at room temperature (**Table 2**). Colonies then spent an additional two weeks in a second quarantine under weekly monitoring against diseases and parasites, before they were transferred to the main system where they proliferated actively (**Fig. 4A, Movie 1**). Because of the high phenotypic variability in *B. diegensis* (Viard et al., 2019), and in the absence of an established genetic marker to assess the genomic diversity of our colonies, we followed the most conservative approach and defined one new strain for each wild isolate. The lineage of all strains, their clonal offspring, lifespan, position in the system and treatments were recorded in an online database with a unique alphanumeric identifier (e.g. strain 1 clone 2: S001C002, strain 131 clone 89: S131C089). Approximately 50 % of the collected specimens could be transferred successfully to our recirculating system.

**Figure 4.**
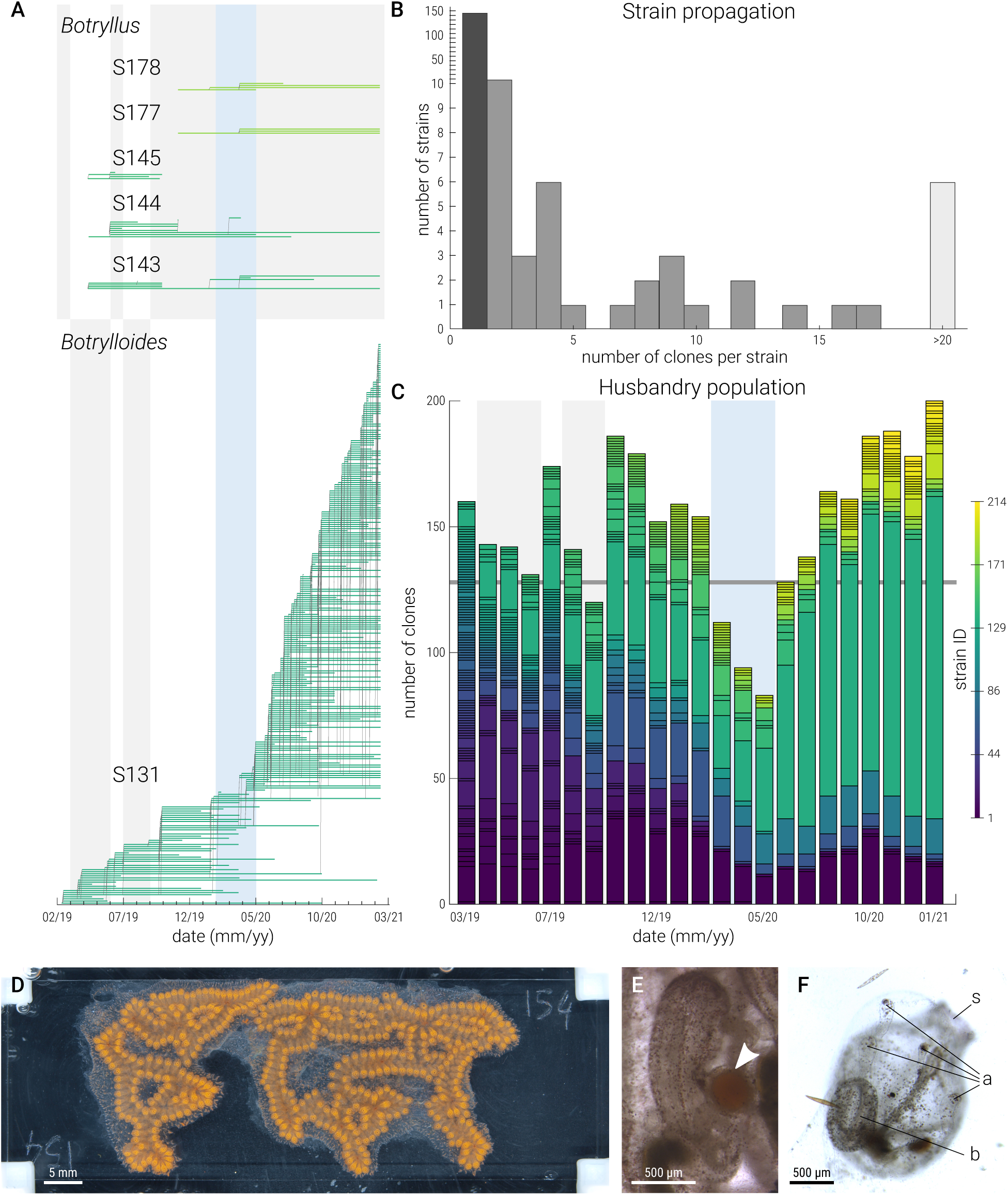
Breeding of colonial ascidians. (**A**) Lineage of representative strains for *Botryllus schlosseri* (S143, S144, S145, S177, S178) and for *B. diegensis* (S131 clone 001 to clone 276). Shown are their propagation by subcloning (from February 2019 till March 2021) as well as the lifespan of individual animals. Major recession events are pointed at by vertical shaded areas (grey and blue), the one concurrent with a sanitary confinement is highlighted in blue. (**B**) Distribution of the number of subclones obtained per strain. The number of strains is depicted with grey bars, the large number of non-proliferating strains is highlighted in dark while the small number of highly proliferative ones in light. Note that the y-axis is not linear above 10. (**C**) Evolution of the number of clones bred in our system shown from March 2019 till February 2021, color-coded by strain ID. Each column represents the number of clones per strain identification number. Major recession events are depicted as in **A**. (**D**) Picture of a relatively large colony composed of over 300 zooids, which became gravid in 2019. (**E**) Picture of a gravid bud with its oocyte visible (arrowhead). (**F**) Picture of a young oozoid recently metamorphosed. Visible organs including its oral siphon (s), ampullae (a) and bud (b).

Over the 3 years of this experiment, we transferred around 100 strains of colonial ascidians, and of *Botrylloides diegensis* in particular, from their natural environment to our recirculating system. We repeatedly observed that some strains declined in the first few months after their acclimatization and ultimately went extinguished without any apparent reason, while neighboring slides with different animals propagated totally fine. Over 75 % of the transferred strains did not propagate sufficiently to be amenable for subcloning (**Fig. 4B**). We could not find any pattern in the premature disappearance of these isolates. However, some strains have been much more productive than others (**Fig. 4C**) and vastly compensated these losses. The proliferating strains reproduced asexually with a blastogenic cycle slightly faster than 7 days (**Fig. 1C**), in our husbandry system tempered at 22 °C. Currently, 6 strains have ended up constituting over 95 % of all the colonies present in our aquaculture (**Fig. 4B**). Although we could have promoted a higher genetic diversity in our lab, this was not an objective in this study. We rather favored fast propagating animals to identify candidate reference strains for in-lab studies where biological replicates and large sample numbers are important. Including the strains created to follow colonies from an undetermined origin, most typically after detaching from their glass slide following subcloning, we have monitored 214 strains for a total of 899 clones. In addition, five strains of *Botryllus schlosseri* bred for research purpose were introduced in the system to assess the suitability of our system for their culture in ASW (**Fig. 4A**). One strain declined after 5.5 months while the other four were bred continuously since their introduction in our recirculating system. Although these colonies showed relatively low proliferation compared to our *Botrylloides*, we could successfully maintain them over prolonged times (15 and 21 months in February 2021, respectively; **Fig. 4A**) undergoing a typical and regular weekly blastogenic cycle (**Supplementary Figure S5**).

In our system, colonies are cultured on numbered microscopy glass slides hanging vertically in the water, which reproduces somewhat their natural environment where we mainly collected them from the lateral sides of floating pontoons. This breeding method has multiple advantages, first for microscopy and general handling of the animals in the lab, second for the ease of use of the badge gripper and the total absence of metal parts (**Fig. 3D**), and finally for the identification and localization of specific colonies within the system. Each slide hangs onto a hook placed onto a PVC tube, itself fixed in the top part of the specimen tanks. Given the space constrains of our system to allow manipulation of each animal individually, each specimen tank can contain up to 32 slides (**Fig. 3C**), for a maximal simultaneous occupancy of 128 slides.

Colonies are checked upon at least once every two weeks, and often daily for slides participating in specific studies. In this occasion, colonies are taken out of the system, cleaned and subcloned if necessary. Subcloning was undertaken for three main reasons. First, for maintenance to compensate the propagation of the animals away from the center of the slide, because this impedes its ease of manipulation and observation. Second to produce clonal biological replicates for experiments. Third, to fuse smaller clones from the same strain to obtain bigger samples for studies where large amounts of tissue were required.

Excitingly, several *B. diegensis* strains have been observed to become gravid (**Fig. 4D-E**). Some of these oocytes even got fertilized as we could identify oozoids developing later on in our system (**Fig. 4F**). Although this result was neither expected nor aimed for, it confirms that our colonies adapted well to our recirculating system and it shows that induction of sexual reproduction is possible under controlled artificial conditions. The environmental conditions required to robustly induce sexual reproduction in *B. diegensis* in our system remain to be determined.

### 3.4. Differential breeding of colonial tunicates

Animals in our recirculating system were bred under continuous feeding following preliminary indications (Blanchoud, unpublished) that such delivery was promoting growth. In addition, this type of feeding has two advantages, first a reduced contamination of the colonies and their slides because of the overall lower concentration of food present in the water, and second a lower daily maintenance as food needs to be prepared only once for multiple days. Continuous feeding was implemented using electronically controlled valves (**Supplementary File 1**). We optimized the frequency and duration of the opening of each valve to reach uniform amounts between all four specimen tanks. On average, valves opened 1.5 s every 3 min, thus delivering around 2 L of diluted food per day. Food was prepared every 4 days to guarantee food quality after culture in our feeding tanks and concentration was adapted to minimize clotting.

Animals were fed with a custom food mix composed of equal parts of two commercial blends of concentrated micro-algae (Shellfish Diet 1800, RotiGrow Plus), supplemented with live rotifers (cultured in-lab), enriched with 1 % (v/v) baker’s yeast and 2 % (v/v) of planktonic supplement rich in amino acids, unsaturated fatty acids and vitamins. To monitor the well-being of our animals, three slides representative of the same three strains were imaged per tank once a week. We manually counted the number of zooids in each picture to estimate the growth rate of our animals and the impact of the different breeding conditions (**Fig. 5A**).

**Figure 5.**
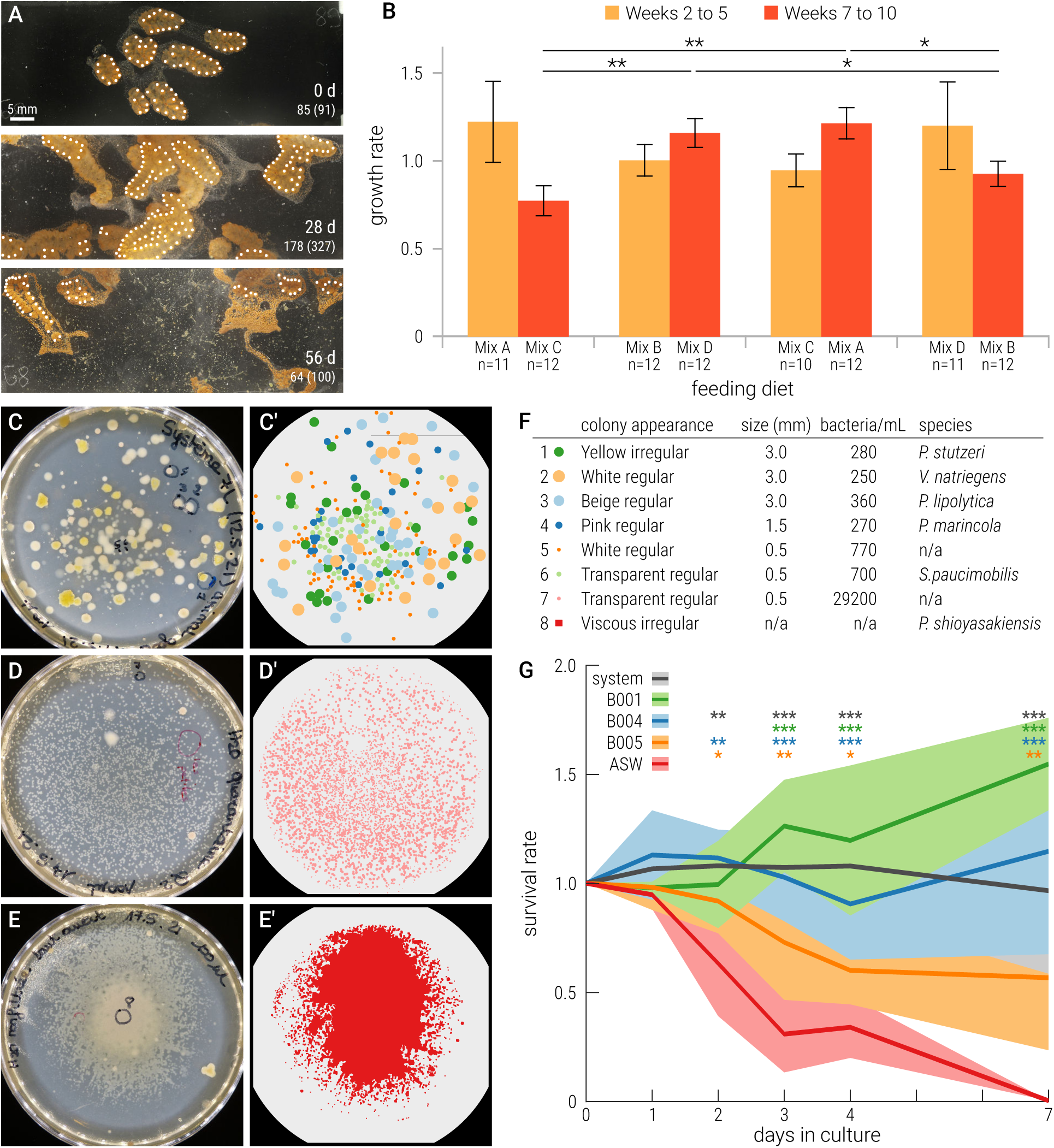
Impact of environmental conditions on the development of *B. diegensis*. (**A**) Sample pictures taken on the specified days during our differential feeding experiment (day 0, 28 and 56). Depicted pictures correspond to the same colony (slide 89) fed with the combination Mix A / Mix B, overlaid with white dots indicating the position of the identified zooids on the front side of the glass slide, the number of identified zooids and the total number of zooids on the slide in parenthesis. (**B**) Growth rates of the colonies in each feeding condition over the indicated number of weeks. Each pair of bars corresponds to feeding conditions that were tested consecutively in the same tank (from left to right: tank 1, tank 2, tank 3, tank 4). Note that transition weeks 1 and 6 were removed from the analysis. Three colonies were monitored weekly in each tank. (**C**) LB agar plate with the bacterial colonies cultured from 100 µL of water from the recirculating system as well as (**C’**) their resulting identification and counting. (**D-D’**) Plate and identification of bacteria from the quarantine. (**E-E’**) Plate and content of an ASW sample. (**F**) Bacterial colonies, appearance, size, *in situ* concentration and tentative species identification. (**G**) Survival rate of colonies (B001 n=3, other conditions n=6) isolated in a 10 L aquaria filled with the water specified, typically ASW inoculated with 50 µL of a bacterial strain. The time points indicated below the x-axis correspond to the quantified times. Statistical significance was measured using Student’s two-tailed *t*-tests compared to the survival in uninoculated seawater and are depicted with * P < 0.05; ** P < 0.01; *** P < 0.001.

Such approach could be used to test the impact of environmental conditions on specific aspects of the specimens’ development, and to adapt our culturing conditions to different species for their breeding. Herein we show one example of such experiments. Animals were fed with four different food mixes: A) 1.6 mL/day ShellFish Diet, 1.6 mL/day RotiGreen Plus, B) 1.1 mL/day ShellFish Diet, 1.1 mL/day RotiGreen Plus, C) 3.2 mL/day RotiGreen Plus, D) 3.2 mL/day Shellfish Diet. All food mixes were complemented with baker’s yeast and planktonic substitute at the same proportion. Animals were bred under these conditions for five consecutive weeks while being monitored weekly. To compensate for any possible bias in specific specimen tanks, we swapped the feeding diets after five weeks, and monitored the same colonies for another five weeks.

Animals fed with food mix A and D outperformed those fed with B and C (**Fig. 5B**). Moreover, animals fed with food mix A and D had positive growth rates (> 1), meaning that each generation produces more animals than the previous one thus leading to exponential growth. Although the differences were not significant in the first set of experiments, we measured the same trends in both sets of feeding experiments. This confirms that the impact of tank specific biases, if any, did not significantly influence our measurements. The low breeding measured with mix B indicates less food than our baseline was not sufficient to sustain a positive growth rate. This confirms that we are not overfeeding our animals. The low breeding measured with mix C indicated that feeding on a blend of two type of algae mix was more beneficial as previously published (Rinkevich and Shapira, 1998). However, the good performance of feeding with D, also a single algae mix, suggests that this food source (Shellfish Diet 1800) is the source of most growth in our species. Overall, this experiment exemplifies how differential breeding can be used in our in-land aquaculture system to study how feeding influences the growth of our colonial ascidians.

### 3.5. A suitable marine microbiota is necessary to the survival of *B. diegensis*

We repeatedly observed that colonies isolated in smaller tanks will present signs of stress and enter into hibernation within a few days, as observed following environmental stresses like low temperatures (Hyams et al., 2017). Based on the communicated crucial role of inoculating artificial seawater with a suitable microbiota (Oliveri, personal communication, unpublished), we decided to investigate the role of bacteria for the survival of *Botrylloides* in ASW. We visually compared the composition of the bacterial community between the water of our recirculating aquarium (**Fig. 5C-C’**), the water of our quarantine one week after its routine replacement (**Fig. 5D-D’**), i.e. without a specific inoculation procedure but in contact with colonies, and ASW left to settle without any contact with animals (**Fig. 5E-E’**). We observed a very clear difference in the diversity of the various communities, with the recirculating system being the only one populated with several types of bacteria (**Fig. 5C-E**). Out of the 8 different types of colonies that we could identify, six were successfully genotyped to five different genera (*Pseudomonas*, *Pseudoalteromonas*, *Psychrobacter*, *Sphingomonas* and *Vibrio*, **Supplementary File 2**), all present in comparable concentrations in the recirculating ASW (**Fig. 5F**).

To determine whether some of these bacteria present in our system participate in the well-being of our ascidians, we measured the short-term survival of colonies in differently inoculated ASW environments (**Fig. 5G**). Out of the six observed types of bacterial colonies, we successfully amplified three (B001, B004, B005) in liquid cultures suitable for inoculation experiments. We measured a very fast decline in colonies bred in uninoculated ASW, with more than half of the zooids (31 ± 18 %) being resorbed within just three days, following an atypical takeover during which a significant fraction of the new generation of zooids was resorbed (**Movie 2**), and all of them being absent after one week (**Fig. 5G**). In contrast, colonies cultured in water from our system maintained their zooids throughout the week. To our surprise, inoculating with 50 µL of single bacterial types was sufficient to significantly improve the short-term survival of *Botrylloides diegensis* in ASW (**Fig. 5G**). B001 (*Pseudomonas stutzeri*) and B004 (*Psychrobacter marincola*) yielded survival rates comparable to those obtained with system water. B005 (undetermined species) led to lower survival closer to those in uninoculated ASW until day 4 although with a reduced decline and a significantly higher survival after seven days (56 ± 33 %, **Fig. 5G**). These results highlight the necessary role of a suitable microbiota in the ASW used for the culture of our colonial ascidians. How and which bacteria participate in the development of tunicates will be an exciting topic to explore.

## 4. Discussion & Conclusions

We here present a hybrid flexible recirculating husbandry setup for the long-term in-lab culture of colonial ascidians. We developed this prototype to sustain our research on the exceptional capacity of *Botrylloides* to undergo whole-body regeneration (Blanchoud et al., 2018a). Yet, in this endeavor we also set out to establish a system that would be suitable for promoting as well as popularizing these colonial ascidians as model organisms more globally. Our achievements in constantly breeding *Botrylloides* over more than 3 years shows that this is indeed possible, and we do hope that it will initiate a renewed interest in studying these fascinating animals in novel places, including in non-coastal ones.

As pioneers in the in-land breeding of ascidians without previous experience with saltwater aquaculture, we obviously initially came across some difficulties with the maintenance of the quality of our recirculating water. Accumulation of noxious ions is the main challenge for such a system. Nitrates in particular tend to accumulate quickly and for the long term because the anaerobic bacteria, that can process this ion, develop much more slowly than the others involved in the rest of the nitrogen cycle (Arrigo, 2005). Although our nitrate filter solved this issue relatively well, the most common solution to this accumulation remains to increase water changes (Zebcare, personal communications). Consequently, the main strategies available to maintain the quality of our ASW were to slowly vary the biological load of our system (i.e. animals and feeding), to use periodic treatments with removable media (e.g. phosphate binding resin), and to perform large water changes (50 % of the total water amount) when needed. To avoid sudden changes in water quality, we thus focused on reducing the variations in biological load on our system. Indeed, the three major regression events we experienced (**Fig. 4A,C shaded areas**) followed a sudden increase in the number of colonies that we naively combined with an increase in feeding to try supporting these additional animals. We experienced a similar issue three times consecutively because we were initially not monitoring all the necessary parameters of water quality, and thus wrongly believed that we had solved the issue at the root of the decline. Instead, the accumulation was transferred to different noxious ions. The third and more drastic regression (**Fig. 4A,C blue area**) was unfortunately concurrent with a sanitary confinement that prevented us from tending properly to our aquaculture. Consequently, the latent issue lasted longer than the two previous ones. Nevertheless, colonies recovered quickly and repopulated our culture setup very efficiently. Moreover, we resorted only twice over the 3 years to large water changes, confirming that the quality of our water is stable.

An important point in controlling the quality of our ASW was to understand that although a particular food source might be beneficial to the growth of our species, it might be imbalanced with respect to our target water parameter values. For instance, we observed that our food mix has a higher pH than our ideal pH 8.2. Although the important dilution that takes place during feeding renders this increase quite minimal, it similarly accumulated over time to yield to a noticeable increase in the whole system (**Fig. 3G**). Properly balancing the pH of the food mix before feeding it to the specimen tanks solved the problem at its root. Consequently, we advise to measure all the parameters of the food mix being delivered to the animals before starting a new feeding experiment. Feeding using live algae (Joly et al., 2007; Brown et al., 2009; Blanchoud et al., 2017) or using home-made blends of vegetable pastes (Kowarsky et al., 2021) might be more appropriate in the context of an in-lab monoculture as presented here and will be experimented in the future.

Colonies of *Botrylloides diegensis* have been cultured in our laboratory for more than 3 years. We have established an in-lab all-year-round stable and fully monitored system which can be adapted to the needs of other tunicates to further popularize this subphylum. Our experiment with *Botryllus schlosseri* (**Fig. 4A**) has shown that this species can adapt to our recirculating ASW aquaculture, although it grows much slower than *B. diegensis*. Yet, colonies look healthy and they do propagate. One possible explanation for this behavior is that *Botryllus schlosseri* colonies thrive at temperatures lower than 20 °C (Boyd et al., 1986), in warmer water they tend to become gravid and slow down their asexual reproduction. Our system allows to set different water temperatures in different specimen tanks, which could be used to optimize *Botryllus* culture.

Similarly, our system could be adapted to breed other species of colonial ascidians that can be cultured on glass slides. Whether all colonial ascidians can be subcloned onto glass slides remain to be determined, but species reported to grow on glass include the Stolidobranchian *Symplegma viride* (Goodbody, 1961) as well as the Aplousobranchian *Diplosoma macdonaldi* (Goodbody, 1961), *Aplidium multiplicatum* (Karande and Nakauchi, 1981), *Clavelina miniata* (Hirose et al., 1996) and *Didemnum vexillum* (Rinkevich and Fidler, 2014). The taxonomic diversity of these species suggests that the culturing techniques that we have established in our recirculating system could be broadly applicable to colonial ascidians pending the environmental conditions are suitably adapted. In addition, we did a short 4-months proof-of-principle experiment with a single *Ciona robusta* specimen that holds interesting promises, provided some improvements in how solitary animals are handled. Subsequently, solitary species could also be bred in our aquarium to culture for instance the highly regenerative *Polycarpa mytiligera* (Gordon et al., 2021).

Controlling the sexual reproduction of the animals in our system will be essential for the propagation of solitary specimens, as well as for multiple aspects of research on colonial ascidians including embryogenesis and transgenesis. While gravid colonies confirm that it is possible for them to reproduce sexually in an artificial environment, it occurred without any apparent relation to a specific breeding condition. In addition, the open configuration of our specimen tanks together with their important water flow-through is not well adapted to handling spawning of tunicates. Further investigation will need to be performed to adapt our in-land aquaculture with published methods used in other species of colonial ascidians (Fletcher and Forrest, 2011; Gasparini et al., 2015b; Joly et al., 2007) both for the induction of spawning as well as for the retrieval and settling of the motile larvae.

A surprising result from our differential breeding experiment is that only one set of data shows statistically significant effects, while the same trends can be observed in the other one. One potential explanation is that some remnant effect of the previous set of feeding experiment was influencing the monitored colonies (**Supplementary Figure S6**). Indeed, because *Botrylloides* have three generations co-existing constantly within the colony (Berrill, 1947), the number of adults observed in the following generation was determined based on the breeding conditions that were in place when this new generation budded, which means around two weeks in our 22 °C system. Consequently, differential breeding experiments might need to be monitored over periods including two additional weeks for buffering between experiments. Alternatively, incorporating only the data from the last week of the experiment could remove this issue, but more colonies would need to be monitored to yield statistically significant results.

Another unexpected result from our breeding experiments is the essential role of bacteria for the breeding of *Botrylloides* in artificial seawater (**Fig. 5G**). Although that the gut microbiota has an influence in the lifespan of other chordates has been well documented (Smith et al., 2017), that a single type of bacteria could have an influence on the short-term survival of an entire colony was a surprise to us. Whether these bacteria have a direct influence on the animals, through some symbiotic process, or indirectly by improving the quality of the seawater, for instance by degrading noxious ions, remains to be determined. However, colonies used in our experiments were initially culture in our recirculating system and consequently should have an established and functional gut microbiota suggesting that a potential interaction through their digestive system is not the most likely. On the other hand, ammonia and other members of the nitrogen cycle have been measured to accumulated quickly in non-filtered seawater (e.g. in our quarantine), possibly degrading the quality of the environment within a few days in small aquaria such as those used in our microbiota experiments. Whether the short-term positive effect of B001 and B004 can be extended to longer time scales will need to be determined, but these results are a first step towards characterizing the crucial role of a suitable microbiota in the success of our recirculating husbandry system.

Overall, we show that *Botrylloides diegensis* can be proficiently bred in-land, that other species such as *Botryllus schlosseri* can also develop in recirculating artificial seawater, and that our system for controlled breeding can provide the necessary platform for the worldwide spreading of colonial ascidians as model organisms.

## 5. Acknowledgments

We would like to thank Dr. Megan Wilson for sharing her experience with breeding *Botrylloides*, Pr. Miles Lamar for providing space for our trials when setting up a flow-through system at the University of Otago (NZ), Pr. Loriano Ballarin for his extensive experience on breeding *Botryllus* in closed systems, Dr. Fabio Gasparini for helping us with the collection of our wild isolates, all the members of these laboratories for useful discussions on the breeding of colonial ascidians, Dr. Stefano Tiozzo for helping with the collection and breeding of colonies in France, as well as Laura Bugada, Margaux de Raemy, Viviane Perraudin and Nathalie Weber for participating to the monitoring of the system.

We thank Dr. Floriane L’Haridon and Pr. Laure Weisskopf for their help in the isolation, characterization, sequencing and amplification of the various microbiota. We thank warmly Pr. Paola Oliveri for her advice to use wild mussels as inoculation media for seawater aquaria. We thank the people from our institute’s workshops for helping us, teaching us and showing us how to modify, improve and develop an aquaculture system. In particular, Ruben Pooley, Alain Werro, Jean-Daniel Niederhauser and François Zosso.

We also thank Mohamed Abshir Mohamed, Jeanne Bruelhart, Lionel Page and Silvia Moreno Forero for their help in taking care of the husbandry setup, and Aude Blanchoud for comments on the manuscript.

Topological map used in Fig. 1B was adapted from the work “Europe laea topography” of Dbachmann and Alexrk2 from Wikimedia Commons. Icons used in Fig. 2 were adapted from work from The Noun Project.

## 6. Author contributions

Simon Blanchoud: Conceptualization, Funding acquisition, Project administration Visualizaton. Marta Wawrzyniak: Writing-original draft. Marta Wawrzyniak, Lluis Matas Serrato, Simon Blanchoud: Investigation, Methodology, Validation, Writing-review & editing.

## 7. Funding

This work was funded by the Swiss National Science Foundation (SNF) [grant number PZ00P3_173981]. The SNF had no involvement in the conduct and preparation of this work.

## 8. Declaration of Interests

The authors have declared no conflict of interest.

## 10. Figure legends

**Movie 1. In-land breeding of *B. diegensis*.** Time-lapse imaging of a colony developing over a period of 25 days.

**Movie 2. Atypical takeover of *B. diegensis* cultured in uninoculated artificial seawater.** Time-lapse imaging of a colony developing in uninoculated artificial seawater (ASW). Recording starts when the colony was transferred from our recirculating system to the uninoculated ASW and runs until an atypical takeover is completed.

**Supplementary Figure S1. Detailed structure of the sump unit.** (**A**) Moving-bed filter, (**B**) refugium and (**C**) main sedimentation tank composing the sump of our husbandry system. (**D**) Fleece filter separated from its underlying moving-bed unit shown in **A**. (**E**) Components of the ceramic beads filtration unit visible in **B**. (**F**) Area between the sump and the chiller where the main circulation pump, the nitrate filter and the leak detection probe are located. The main components visible in the panels are identified using their ID as listed in Table 1.

**Supplementary Figure S2. Detailed structure of the feeding unit.** (**A**) Feeding tank connected to the delivery valve for dispensing food in the specimen tank located underneath. Each feeding tank is composed of (**B**) a 10 L cylindrical container, (**C**) a micro-porous air diffuser and (**D**) a lid with a clear compartment for hosting the lighting bars. Components ID are specified in Table 1.

**Supplementary Figure S3. Detailed structure of the controlling units.** (**A**) Main control panel with the Apex controller and its various modules. (**B**) Behind the husbandry system where the solenoid valve controls the input of deionized water for topping-up the sump. (**C**) Components of the custom micro-controller unit for continuous feeding of our colonies. (**D**) Lights and feeding control units are installed behind the feeding tanks. Components ID are specified in Table 1.

**Supplementary Figure S4. Detailed structure of the air and salts units.** (**A**) The main air pump is isolated from the husbandry system and connects to a custom manifold. The pH unit is located on the left side of the system. (**B**) The manual control valve for the pH buffer is accessible from behind the main system. (**C**) The brine tank is located on the right side of the system and is pumped automatically by the two peristaltic pumps visible in **A**. Components ID are specified in Table 1.

**Supplementary Figure S5. *Botryllus schlosseri* in our system.** Asexual reproduction cycle in our breeding system. The colony growing on slide 75 is shown as an example of the observed weekly blastogenic cycle (days 0 to 7). Similar to *Botrylloides diegensis* (Fig. 1C), blastogenesis starts and ends with the takeover stage. White square highlights the system that is magnified and followed every day throughout the week, overlaid with the corresponding number of days after takeover.

**Supplementary Figure S6. Average growth rates per specimen tank.** Average (bold line) and standard deviation (shaded area) of the growth rates of the three colonies monitored in each specimen tank throughout the feeding experiments. Color-codes represent to the feeding condition depicted in Fig.5B, with the gray section depicting the transitory week. Growth rates are overlaid with the corresponding feeding mix and specimen tank.

**Supplementary File 1. Design and control of our custom microcontroller for continuous feeding.** This archive contains all the documents required to reproduce our microcontroller. The file used to design and produce the custom PCB board (controller_design.fzz) using Fritzing, an Open Source software. A schematic representation of that PCB board (controller_design_scheme.pdf) and two vectorized visualizations of the PCB connections, one for each side of the board (controller_design_pcb_top.pdf, controller_design_pcb_bottom.pdf). A list of the electronic components required to assemble the PCB board (controller_components.txt) and the MicroPython code used to control the valves (controller_code.py).

**Supplementary File 2.** Sequences used for DNA barcoding, PCR primers and resulting species determination.

## Notes

### Competing Interest Statement

The authors have declared no competing interest.

### Summary of Updates

This manuscript is the version accepted for publication at Developmental Biology, following two round of peer-review. Additional data and multiple clarifications were added to the article, at all level of the manuscript.

