## Supplementary figures and images for "Artificial seawater based long-term culture of colonial ascidians"

### controller_design_pcb_bottom.pdf

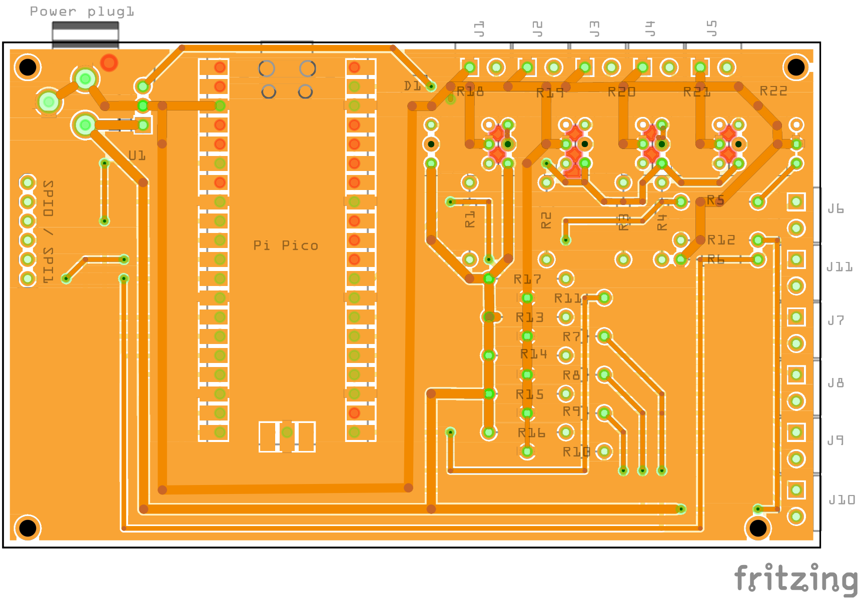

### controller_design_pcb_top.pdf

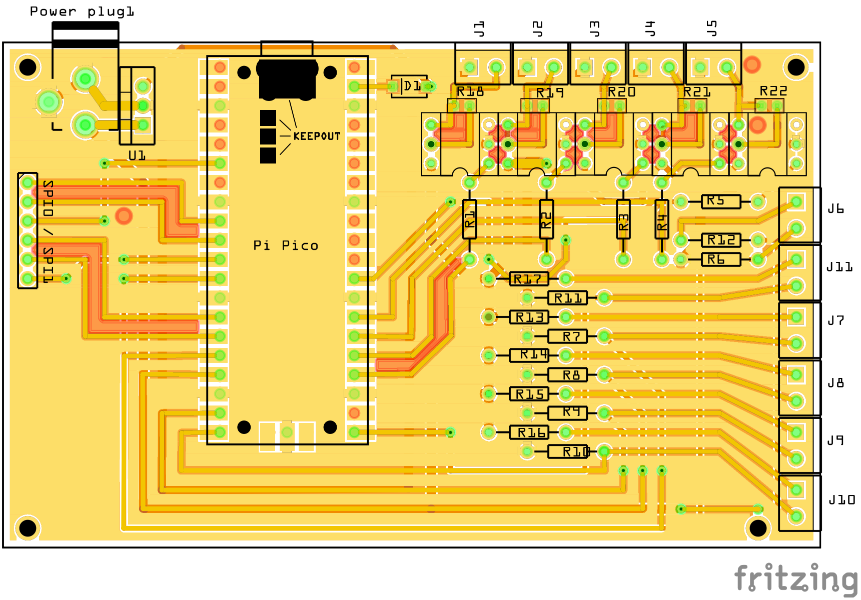

### controller_design_scheme.pdf

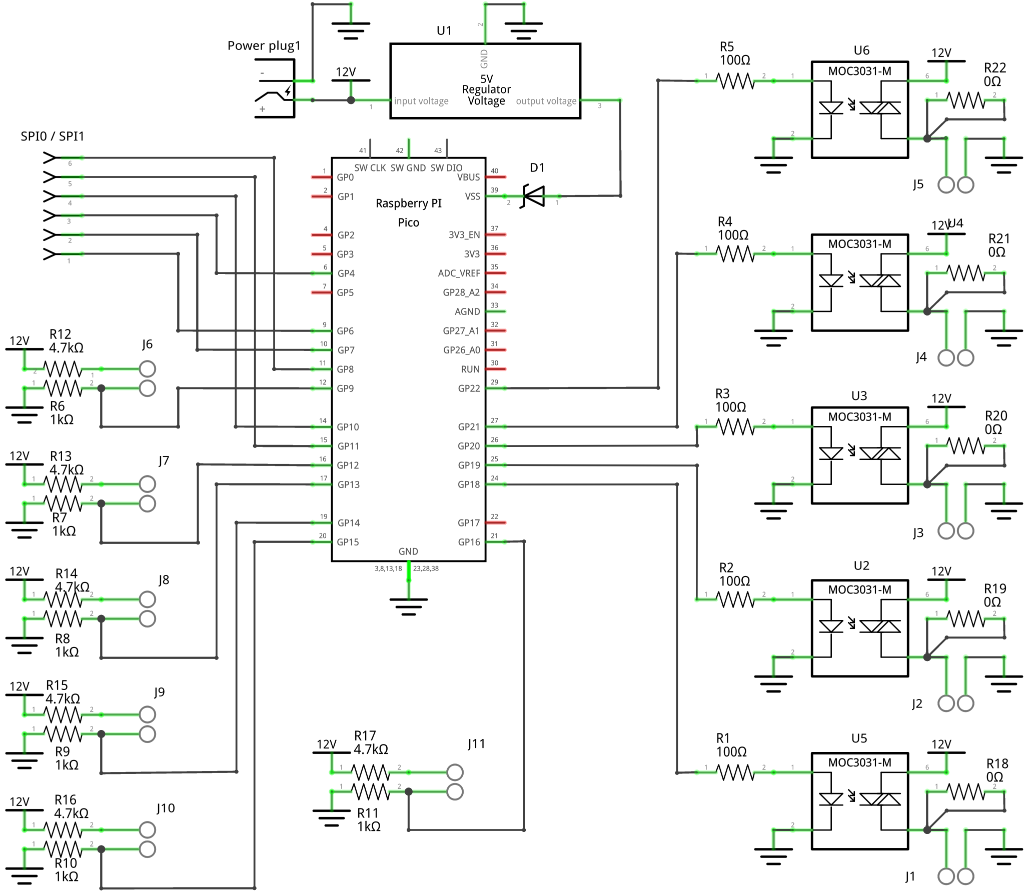

### Supplementary Figure S1

#### Supplementary Figure S1

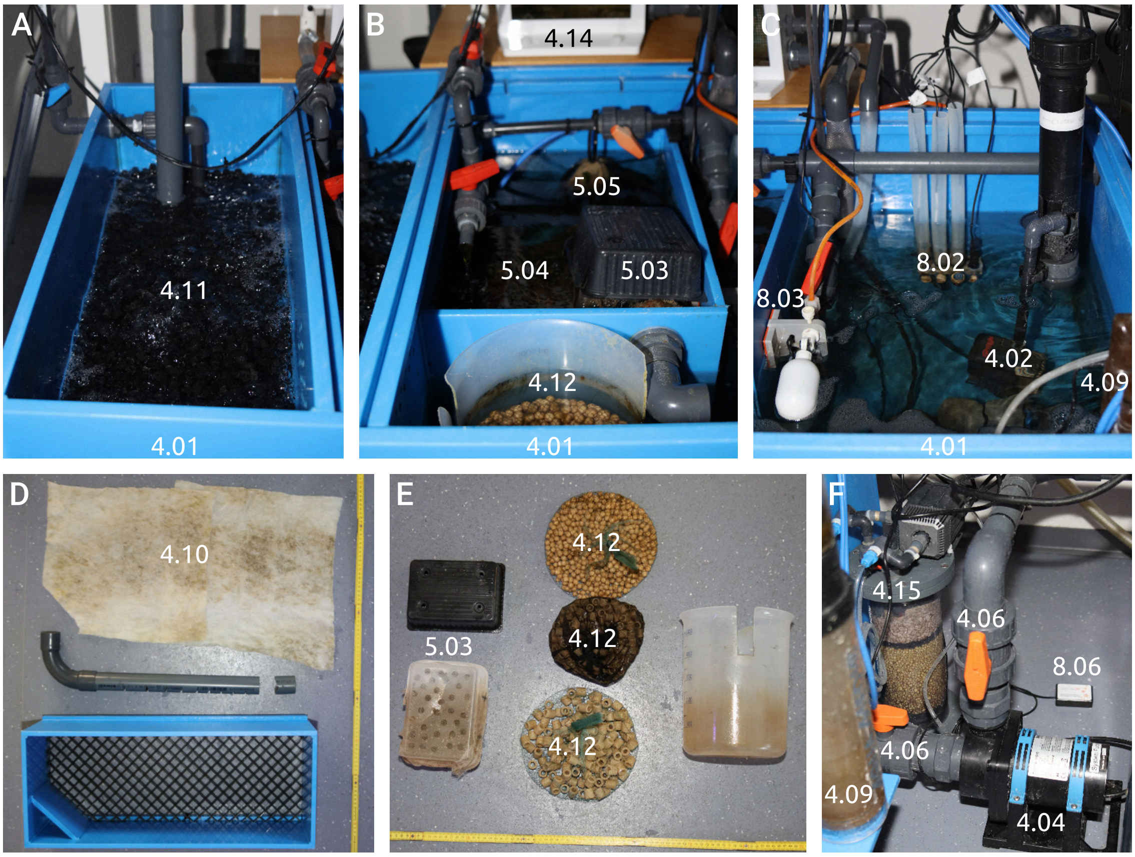

### Supplementary Figure S2

Supplementary Figure S2

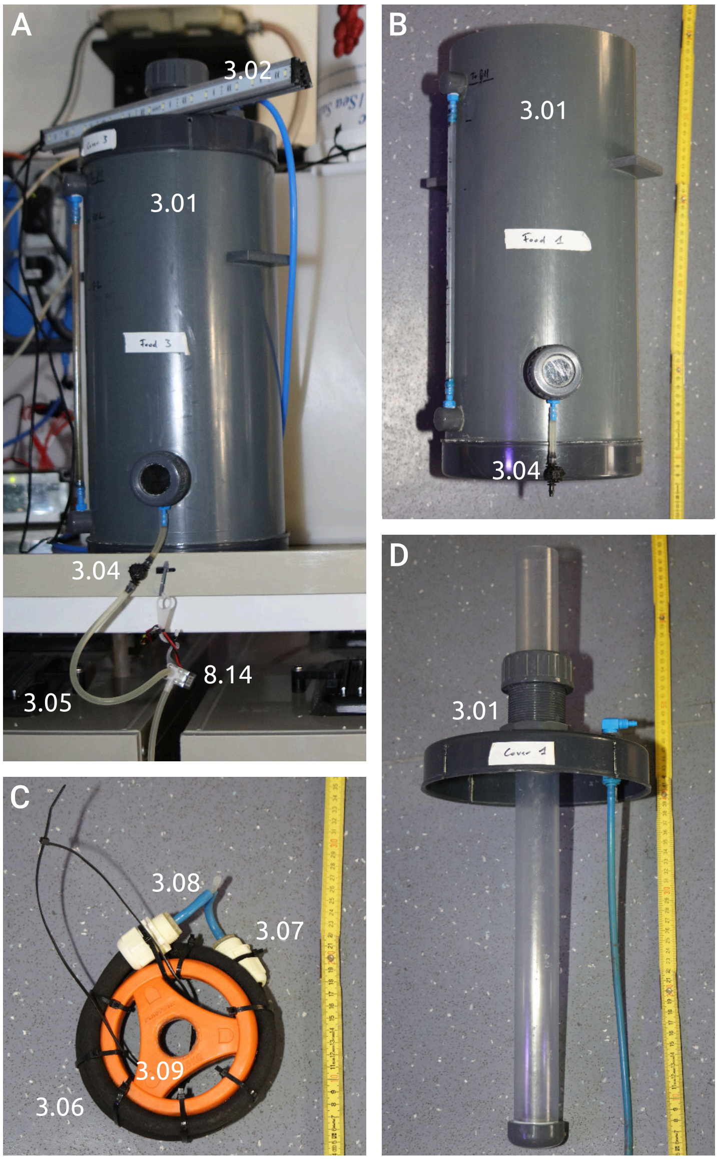

### Supplementary Figure S3

Supplementary Figure S3

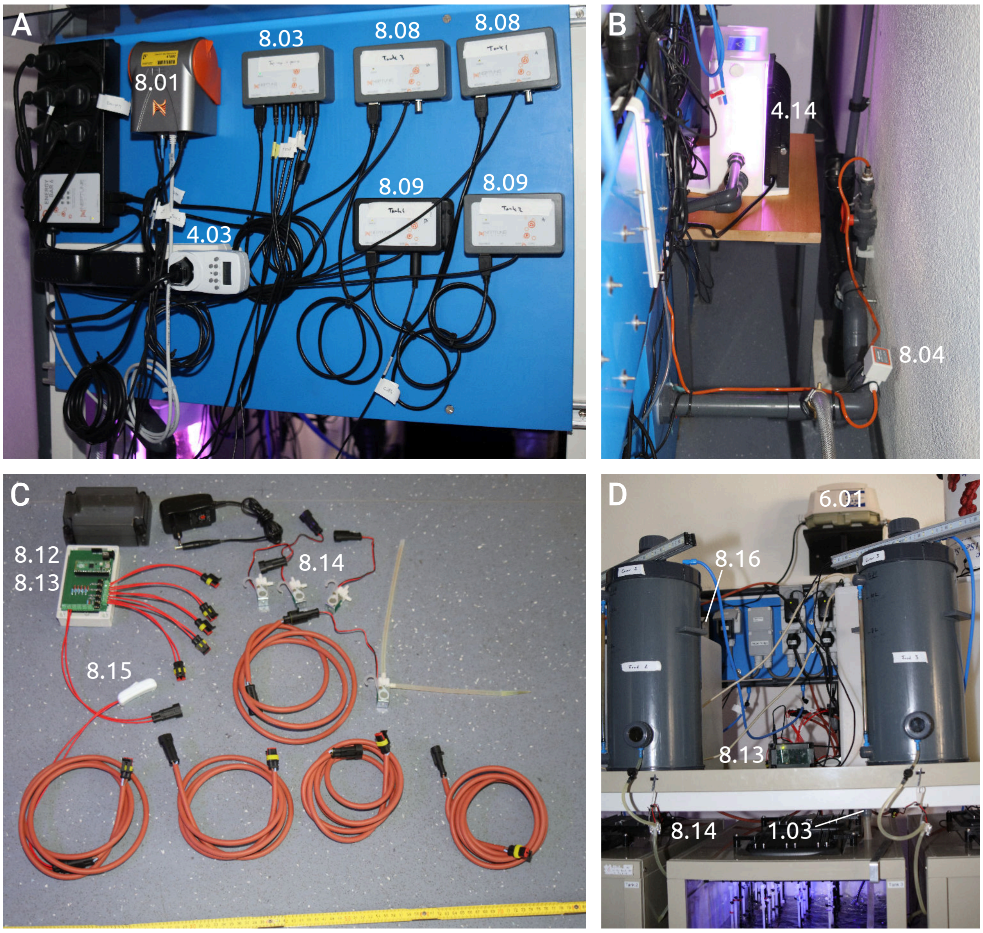

### Supplementary Figure S4

#### Supplementary Figure S4

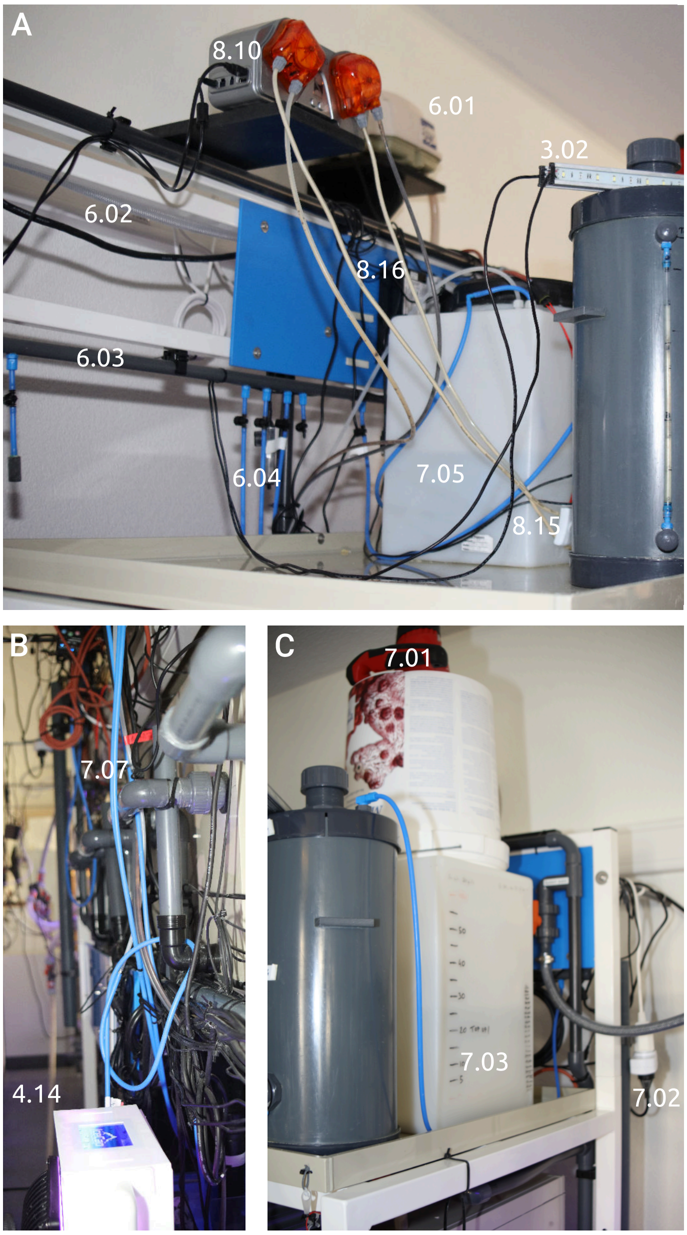

### Supplementary Figure S5

##### Supplementary Figure S5

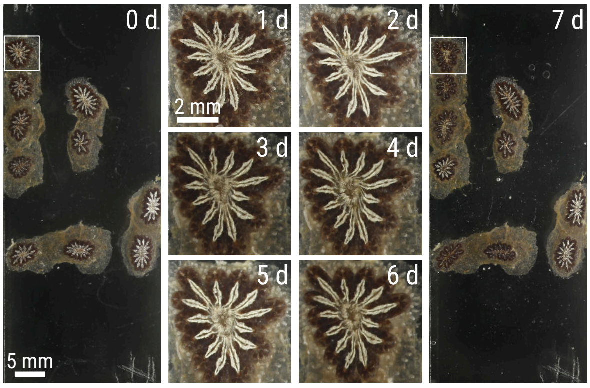

### Supplementary Figure S6

##### Supplementary Figure S6

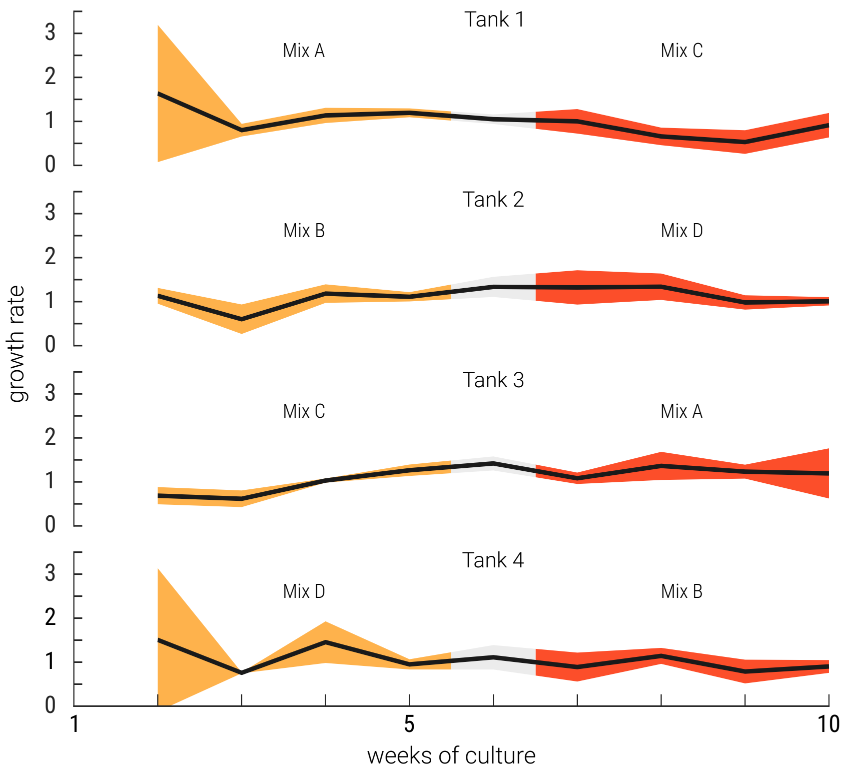
