## Supplementary File 1 for "Artificial seawater based long-term culture of colonial ascidians": controller_code.py

```
from machine import Pin, Timer
import time

# Pin assignment for IN/OUT
pins_out = [18, 19, 20, 21, 22]
pins_in = [9, 12, 13, 14, 15, 16]

# Main led
led = Pin(25, Pin.OUT)

# Interval between deliveries in ms and the corresponding timer
loop = 160000
tim1 = Timer()
tim2 = Timer()

# Duration of delivery for each valve in ms
timings = [1540, 1540, 1540, 1540, 1540]
mtime = max(timings)

# Array for the various objects to be created
timers = []
valves = []
inputs = []
flags = []

# Global variables used to share information between the functions
state = 0
reset = 0
increment = 0

# Assign the valves to the pins and initialize
# the corresponding timers
for idx, pin in enumerate(pins_out):
    valves.append(Pin(pin, Pin.OUT))
    valves[idx](False)
    timers.append(Timer())

# Assign the potential inputs
for idx, pin in enumerate(pins_in):
    inputs.append(Pin(pin, Pin.IN, Pin.PULL_DOWN))
    flags.append(0)

# Callback for each valve at the end of its delivery time
def delivery(alarm):
    # Loop through all timers
    for idx, tim in enumerate(timers):
        # Find which one ended
        if timers[idx]==alarm:
            # Turn off the corresponding valve
            valves[idx].toggle()

# Main callback, starts food delivery and initiates the timers for closing them
def tick(timer):
    # Retrieves the global variables
    global increment, state, reset
    
    # Counter used to compensate for the loss of pressure
    increment = increment + 1
        
    # Computes that actual amount of compensation needed
    dt = float(increment)/25;
    if (dt<0):
        dt = 0
    elif dt>mtime:
        dt = mtime 
    scale = 1 + dt/mtime
    
    # State 0 is constantly opened valves
    if state==0:
        
        # Do not reset straightaway the compensation in case it was a burst
        reset = reset + 1
        
        # Once we counted enough of them, do reset the compensation
        if reset > 10:
            increment = 0
     
    # Otherwise we deliver the food as expected
    else:
        # Loop through all the timings
        for idx, dur in enumerate(timings):
            # If necessary, start a timer with the corresponding duration and open the valve   
            if dur>0:
                timers[idx].init(callback=delivery, mode=Timer.ONE_SHOT, freq=(1000/(scale*dur)))
                valves[idx].toggle()
                time.sleep(0.02)

# Secondary loop to monitor the inputs. Interrupts could be used but seem to take a lot of ressources.
def interrupts(timer):
    # Retrieve the global variables
    global state, reset
    
    # Blinking specials for state 0
    if state==0:
        # Visually shows that we changed state
        led.toggle()
    
    # Loop through all the inputs
    for idx, pin in enumerate(inputs):
        
        # Check the value of each pin, buffer its value
        count = 0
        for i in range(10):
            count += pin.value()
            time.sleep(0.001)
        
        val = (count > 5)
        
        # If it changed since booting time, something happened
        if flags[idx] != val:
            
            # Store the change and adapt the state of the program
            flags[idx] = val
            state = val
            
            # We just registered a switch to state 0
            if val == 0:             
                # Start counting for increment reset
                reset = 0
                
                # Stop all the timers
                for tim in timers:
                    tim.deinit()
                    
                # Open all valves
                for valve in valves:
                    valve(True)
                    
            # This is back to state 1
            else:
                # Visually shows that we are in state open
                led.value(val)
                
                # Close all valves, delivery will be reinitiated by the main loop
                for valve in valves:
                    valve(False)

# Start the led
led.value(state)

# Start the main loop
tim1.init(freq=(1000/loop), mode=Timer.PERIODIC, callback=tick)

# Start the secondary interrupt loop
tim2.init(freq=1, mode=Timer.PERIODIC, callback=interrupts)
```
